# ER Stress-Induced β-Cell Apoptosis is Linked to Novel Select Lipid Signaling at the Transcriptional Level: Implications in T1D Developmen*t*

**DOI:** 10.64898/2026.03.02.708596

**Authors:** Xiaoyong Lei, Anil K. Challa, Susan E. Nozell, Tomader Ali, Daniel J. Stephenson, Andrew Nafzinger, Chad S. Hunter, Adam R. Wende, Ernesto S. Nakayasu, Ying Gai-Tusing, Charles E. Chalfant, Sasanka Ramanadham

**Author notes:** **<u>Corresponding Author</u>** Sasanka Ramanadham, PhD, Professor, Department of Cell, Developmental, and Integrative Biology (CDIB), Senior Scientist, Comprehensive Diabetes Center, University of Alabama at Birmingham (UAB), Shelby Biomedical Research Building, Rm. 1205, Birmingham, AL 35294-2182, USA.

## Abstract

Type 1 diabetes (T1D) is a consequence of β-cell death. ER stress precedes T1D onset and prolonged ER stress in β-cells can lead to β-cell apoptosis. We reported that lipid signaling generated by the Ca^2+^-independent phospholipase A_2_β (iPLA_2_β), encoded by *Pla2g6*, participates in ER stress-mediated β-cell apoptosis. β-Cell membranes are enriched in arachidonic acid containing glycerophospholipids and the iPLA_2_β catalyzes the hydrolysis of arachidonic acid in ER stressed β-cells. Metabolism of arachidonic acid leads to the generation of various proinflammatory lipids, raising the possibility that they contribute to ER stress and β-cell death leading to T1D. However, molecular mechanisms by which such β-cell-iPLA_2_β-derived lipid (iDL) signaling contributes to β-cell apoptosis are not understood. It is well known that ER stress-mediated β-cell apoptosis is associated with induction of transcription factors, NFκB and STAT1. We report here that both induce *Pla2g6* and, unexpectedly, we find that iPLA_2_β, which lacks DNA-binding motifs, associates with *NFkB*, *Stat1*, and *Pla2g6* promoter regions. Consistently, p65-NFκB and pSTAT1 induction is reduced with select inhibition or knockdown of iPLA_2_β. Surprisingly, iPLA_2_β expression is also reduced by select inhibition of iPLA_2_β, raising the possibility of feedback regulation by iDLs. In support, we find that select iDLs, recognized to be proinflammatory, enhance association of iPLA_2_β with *Pla2g6*, *Nfkb*, and *Stat1* promoter regions leading to induction of all three gene products and β-cell apoptosis. Our findings reveal previously unrecognized transcriptional regulation by iDL signaling and, iPLA_2_β itself, that leads to gene products that promote β-cell apoptosis. Analogous findings in human islets validate this mechanism raising the possibility that targeting select lipid signaling can reduce ER stress in β-cells and ameliorate T1D development.

## Introduction

Type 1 diabetes (T1D) is a consequence of autoimmune destruction of β-cells and the involvement of immune cells is extensively-studied^1^. The β-cells also generate signals that support immune responses by mechanisms that are not well-understood. Our on-going studies highlight roles of select lipid signaling in β-cell apoptosis^2–9^ and T1D development^10–13^. While several factors have been identified as inducers of β-cell apoptosis, there are no studies linking it with lipid signaling generated by β-cells themselves.

Accumulating evidence suggests that β-cell apoptosis due to ER stress is a critical contributor to β-cell death leading to T1D^14–27^. Importantly, ER stress in β-cells precedes T1D onset in both NOD, a spontaneous autoimmune mouse model of T1D, and humans^28,29^. Despite the mounting evidence linking ER stress with β-cell apoptosis leading to T1D, underlying molecular mechanisms triggered in the β-cells themselves that contribute to their death are not well-understood.

Our work reveals that the group VIA phospholipase A_2_ (iPLA_2_β) participates in β-cell apoptosis^2,4,5,30^. The iPLA_2_β hydrolyzes the *sn*-2 substituent from membrane glycerophospholipids to release fatty acids^31^. In islets, iPLA_2_β is predominantly localized in β-cells^2,4,32^, and is induced in ER-stressed β-cells^2,4,32^. Various strategies (inhibitors, siRNA, and genetic-modulation) indicate a critical role for iPLA_2_β in β-cell apoptosis due to ER stress^2,30,33–37^. We reported that inhibition or reduction of iPLA_2_β in the NOD preserves β-cell mass and glucose tolerance, and reduces T1D incidence^10,13^, supporting a critical role for iPLA_2_β-derived lipids (iDLs) in T1D development.

The β-cell membranes are enriched in membrane glycerophospholipids containing arachidonic acid in the *sn*-2 position. ER stress in β-cells is associated with induction of iPLA_2_β at the transcription level^30^. This can lead to increased hydrolysis of arachidonic acid, which can be metabolized by various enzymes to generate oxidized lipids, many of which are proinflammatory^38^. These include prostaglandins (PGs) that are generated via cyclooxygenases and are linked to stress responses^39–43^. They are also recognized to serve signaling/co-factor functions to promote transcriptional activity^44–47^. However, the underlying molecular mechanisms these iDLs trigger in β-cells to promote their own death have not yet been identified.

Multiple transcription factors are activated in stressed β-cells leading to induction of genes encoding factors which contribute to β-cell apoptosis^48–50^. Among these are NFκB and STAT1 that are induced in β-cells during ER stress^48–50^. Herein, we report that in ER stressed β-cells, NFκB and STAT1 induce *Pla2g6* at the transcription level and intriguingly, that iPLA_2_β, which does not contain DNA-binding motifs, associates with *Nf*κ*b* and *Pla2g6* promoter regions. Our findings suggest a novel function of iDLs as co-factors that facilitate transcriptional regulation by iPLA_2_β to trigger generation of gene products that promote ER stress and β-cell apoptosis. Hence, our study identifies potential candidate lipids that could be targeted to reduce ER stress and β-cell death and consequentially, ameliorate T1D development.

## Methods

### General materials

Coomassie reagents, SDS-PAGE reagents, kaleidoscope pre-stained molecular mass standards, and Triton X-100 (161-0324, 161-0407; BioRad, Hercules, CA); *S*-BEL (iPLA_2_β inhibitor), PGE_2_ (10006801, 14010; Cayman Chemicals, Ann Arbor, MI); acetyl histone H4 (AcH4antibody, normal mouse IgG, protein A beads (06-911, 12-371, 16-125; EMD Millipore, Billerica, MA); Collagenase 2, fetal bovine serum (FBS) and common cell culture reagents (17101-015; Gibco, Carlsbad, CA); Immobilin-P PVDF membrane (IPVH00010; Millipore Corp., Bedford, MA; protease inhibitor cocktail (D4693), alpha-MEM media (P8340), and common reagents (Sigma Chemical Co., St. Louis, MO); and enhanced chemiluminescence reagent, (34095; Thermo-Fisher, Waltham, MA).

### Cell culturing

<u>Rat-derived INS-1 β-cells</u> were cultured in Roswell Park Memorial Institute (RPMI, Cat. 11875093, Thermo Fisher Scientific Inc. <u>Waltham, MA</u>) 1640 medium, containing 11 mM glucose, 10% fetal calf serum, 10 mM HEPES buffer, 2 mM glutamine, 1 mM sodium pyruvate, 50 mM β-mercaptoethanol (BME), and 0.1% (w/v) each of penicillin and streptomycin in cell culture conditions (37°C, 5%CO_2_/95% air). <u>Mouse-derived</u> <u>MIN6 β-cells</u> were cultured in Dulbecco’s modified Eagle’s medium (DMEM, Life Technologies Corporation, Grand Island, NY) containing 15% fetal calf serum, β-mercaptoethanol (150 µM), and 0.1% (w/v) each of penicillin and streptomycin in cell culture conditions (37°C, 5%CO_2_/95% air). For any given experiment, except where indicated, the cells were seeded in 6-well plates and used at ∼70% confluency. Data (Figs. 1 and 3) generated in INS-1 cells were used to guide all the remaining studies in MIN6 cells.

**Figure 1.**
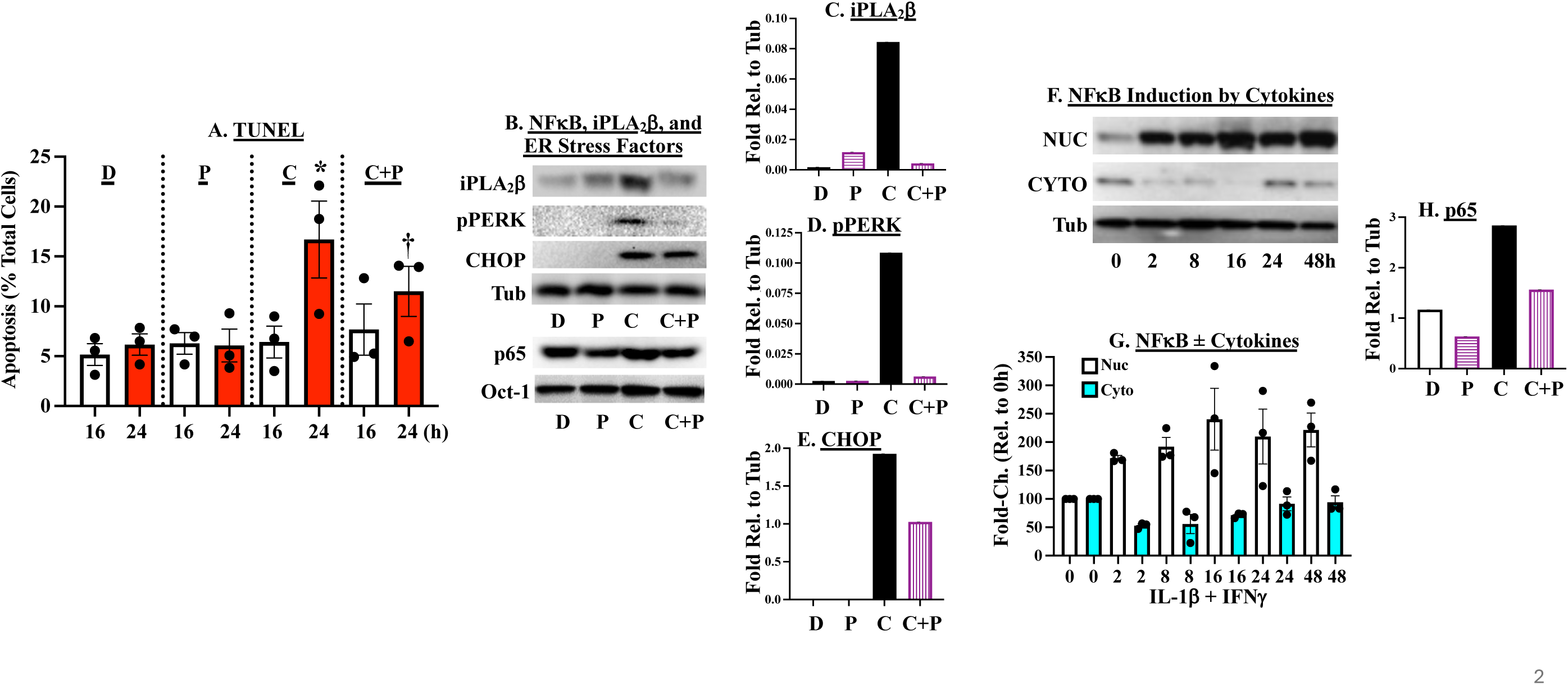
NFκB Activation During ER Stress-Induced β-Cell Apoptosis is Associated with iPLA β Induction. INS-1 β-cells were treated with <u>D</u>MSO or <u>C</u>ytokines (50 U/ml IL-1β + 150 U/ml IFNγ) for 16h or 24h (**A**), 16h (**B-E, H**), or 2-48h (**F-G**). **A**. Cells were processed for TUNEL analyses and mean ± SEMs of %apoptotic cells, relative to total cells, are presented. (^*,†^Significantly different from corresponding DMSO group, p < 0.05; ^†^significantly different from corresponding DMSO and cytokine groups, p < 0.05, n=3/group.) **B-E, H**. Representative blot from cells treated without or with <u>P</u>BA for iPLA_2_β, p65-NFκB, pPERK, and CHOP expression and corresponding densitometries. Tubulin and Oct-1 (nuclear marker) were uses as loading controls in n=3 independent analyses. Cumulative densitometries are presented in **Suppl. Fig. 2. F, G**. Representative blot of iPLA_2_β expression in cytosolic and nuclear fractions prepared, as described^7^, and cumulative densitometry.

### *Pla2g6* knockdown by siRNA

Cells were seeded in 6-well tissue culture plates () in 2 ml antibiotic-free normal growth medium supplemented with FBS. After incubating the cells (37°C, 5%CO_2_/95% air) until the cells 60-80% confluency was reached. The group VI iPLA_2_ siRNA (m) (sc-43820, Santa Cruz, Dallas, TX) duplex solution was then added directly to the dilute Transfection Reagent (sc-36868, Santa Cruz Biotechnology, Dallas, Texas) using a pipette. After gentle mixing by pipetting, the mixture was incubated for 20 min at room temperature in cell culture hood and then transferred to an incubator (37°C, 5%CO_2_/95% air). After 5-7h, 1 ml of normal growth medium containing 2X the normal serum and antibiotics concentration (2x normal growth medium) was added to the transfection mixture. After an additional 18-24h, the medium was aspirated and replaced with fresh 1x normal growth medium prior experimentation.

### *Pla2g6* knockdown by CRISPR-Cas9 protocol

Constructs that encoded guide 0 (control) or guide1 and 2 were used to generate control cell line and *Pla2g6* knockdown cells by using Lipofectamine™ 3000 Transfection Reagent (L3000015, Thermo Fisher Sci). Transfected cells were selected with 0.25 μg/mL puromycin, and single-cell clones expanded and used for treatments.

### Treatments

Cells were treated with DMSO or cytokines (50 U/ml IL-1β + 150 U/ml IFNγ) (401-ML and 485-MI; R & D Systems, Minneapolis, MN) and processed for immunoblotting, TUNEL, ChIP, and lipidomics analyses. To assess the impact of select pathways, cells were pretreated (1h) with inhibitors of iPLA_2_β (*S*-BEL, 1 µM, Cayman Chemicals, Ann Arbor, MI, #10006801 or FKGK18 (5 x 10^-8^M, generously provided by Dr. George Kokotos, University of Athens, Greece); iPLA_2_γ (*R*-BEL, 1 µM, #10006800), cPLA_2_ (Cay10502, 10 µM, #10008657), STAT1 activation (AZD 1480, 10 µM, #10702) from Cayman Chemicals, Ann Arbor, MI,); and NFκB activation (Bay11-7082, 10 µM, Sigma-Aldrich Inc., St. Louis, MO, #196870). To assess the role of select pathways, cells were treated with 10 nM PGE_2_ (#14010), 10 µM PGF_2_α (#16010), 10 µM 8-Iso PGF_2_α (#16350, Cayman Chemical, Ann Arbor, Michigan)

### Protein analyses

The 1° antibodies for targeted proteins included: iPLA_2_β (sc-166616, 1:1,000), p65-NFκB (#sc-372, 1:1,000), CHOP (#sc-575, 1:1,000), IKBα (sc-371, 1:1000), tubulin (#sc-8035, 1:1,000), and Oct 1 (#sc-8024, 1:1,000) from Santa Cruz Biotechnology, Dallas, Texas; and pPERK (Cell Signaling Danvers, MA, #3179, 1:500). As quantitation of p65-NFkB, relative to tubulin and Oct-1 (nuclear marker), was found to be similar, only tubulin was used as loading control in subsequent experiments. The 2° antibodies concentrations were 1:10,000 (Goat anti mouse, sc-2031, Goat anti rabbit, sc-2030, Santa Cruz Biotechnology, Dallas, Texas). Immunoreactive bands were visualized by enhanced chemiluminescence. In the text, unless otherwise noted, representative immunoblots and corresponding densitometry quantitations are presented. The cumulative quantitations ± SEMs are presented as Supplemental figures.

### *In situ* detection of DNA cleavage by TUNEL staining

Cells were harvested and washed twice with ice-cold PBS and processed for TUNEL analyses using In Situ Cell Death Detection Kit (Cat. 1684795, Roche Diagnostic Corp., Indianapolis, IN) The cells were then immobilized on slides by cytospin and fixed with 4% paraformaldehyde in PBS (pH 7.4, 1h, room temperature). The cells were then washed with PBS and incubated in permeabilization solution (0.1% Triton X-100 in 0.1% sodium citrate in phosphate-buffered saline for 30 min at room temperature). The permeabilization solution was then removed and TUNEL reaction mixture (50 μL) added, and the cells incubated (1h at 37°C) in a humidified chamber. The cells were tjen washed again with PBS and counterstained with 1 μg/mL DAPI (4’, 6’-diamidino-2-phenylindole, DAPI Cat. 62248, Thermo Fisher Scientific Inc., Waltham, MA in phosphate-buffered saline for 10 min to identify cellular nuclei. The incidence of apoptosis was assessed under a fluorescence microscope using a FITC filter. Cells with TUNEL-positive nuclei considered apoptotic. DAPI staining was used to determine the total number of cells in a field. A minimum of four fields per slide was used to calculate the percent of cells that were apoptotic.

### ChIP protocol

The ChIP analyses were performed by using Magna ChIP® A – a chromatin immunoprecipitation kit (Sigma-Aldrich Inc., St. Louis, MO, #17-610 Magna ChIP®), as previously described^51,52^. Briefly, MIN6 β-cells (3 x T225 flasks) were cultures in the absence (DMSO only) or presence of treatments for different durations, as described in appropriate results sections. The nuclei from cross-linked cells were resuspended in Tris/EDTA. The soluble chromatin was adjusted in RIPA buffer and pre-cleared with salmon sperm blocked protein A beads (Cat. 16-125; EMD Millipore, Billerica, MA). Immunoprecipitation was performed with 5 μg of antibodies directed against acetylated histone (AcH4), iPLA_2_β or IgG. After pre-clearing and before immunoprecipitation, equal amounts of sonicated DNA (10 % volume of each sample) were reserved for qPCR analysis. The *Pla2g6, Nfkb, Stat1, Gcg, Pepck*, and *Il-6* promoter regions were probed with specific primers (**Table 1**) against the immunoprecipitated DNA by qPCR. Real-time qPCR was performed using Fast SYBR® Green PCR Master Mix (Thermo Fisher Scientific, Auburn, AL, #A25742) and carried out in a plate-based LightCycler® 480 System (Roche Life Sciences), as we described^5,10,13^. AcH4 was used to confirm transcriptional activity and IgG as a negative control.

**Table 1.**
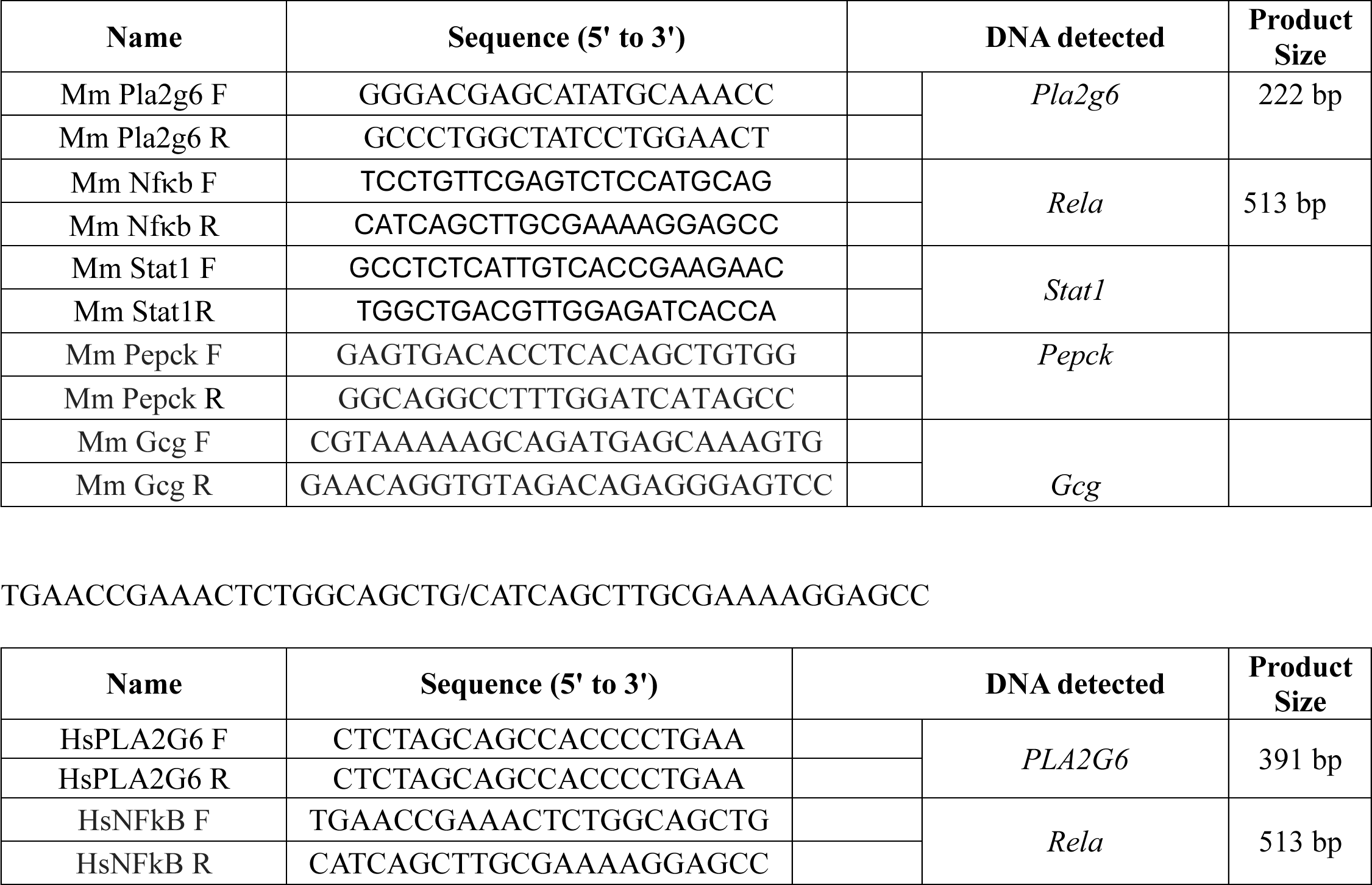
ChIP Primers.

### Human islets

Islets from healthy donors were obtained through Integrated Islet Distribution Program (IIDP) Alberta Islet Distribution Program (AIDP)^53,54^, Clinical islet Laboratory, Univ. Alberta Hospital. The islets upon receipt were immediately cleaned of non-islet material under a microscope. The islets were counted and cultured in RPMI 1640 medium (containing 10% FCS, 200 mM glutamine, 1% of 100 × penicillin-streptomycin, 25 mM HEPES) for 24h at 37°C under an atmosphere of 5%CO_2_-95% air. The islets were then distributed into dishes for treatment and different analyses.

#### Lipidomics

Islets were then treated with cytokines (100 U/ml IL-1β + 300 U/ml IFNγ) from 2 to 36 h. In some experiments the islets were cultured with cytokines ± *S*-BEL at peak response time. The media were collected for analysis of eicosanoids by UPLC ESI-MS/MS analyses, as described^10,12,13^. The media was combined with an IS mixture comprised of 10% methanol (400 μL), glacial acetic acid (20 μL) and internal standard (20 μL) containing the following deuterated eicosanoids (1.5 pmol/μL, 30 pmol total). (All standards purchased from Cayman Chemicals): (d4) 6keto-prostaglandin F_1_α, (d4) prostaglandin F_2_α, (d4) prostaglandin E_2_, (d4) prostaglandin D_2_, (d8) 5-hydroxyeicosatetranoic acid (5-HETE), (d8) 12-hydroxyeicosa-tetranoic acid (12-HETE) (d8) 15-hydroxyeicosa-tetranoic acid (15-HETE), (d6) 20-hydroxyeicosa-tetranoic acid (20-HETE), (d11) 8,9-epoxyeicosatrienoic acid, (d8) 14,15 hpoxyeicosa-trienoic acid, (d8) arachidonic acid, (d5) eicosapentaenoic acid (EET), (d5) docosahexaenoic acid, (d4) prostaglandin A_2_, (d4) leukotriene B_4_, (d4) leukotriene C_4_, (d4) leukotriene D_4_, (d4) leukotriene E_4_, (d5) 5(*S*),6(*R*)-lipoxin A_4_, (d11) 5-iPF_2_α-VI, (d4) 8-iso prostaglandin F_2_α, (d11) (±)14,15-dihydroxyeicosatrienoic acid (DHET), (d11) (±)8,9-DHET, (d11) (±)11,12-DHET, (d4) prostaglandin E_1_, (d4) thromboxane B_2_, (d6) dihomo-gamma linoleic acid, (d5) resolvin D2, (d5) resolvin D1 (RvD1), (d5) maresin2, (d7) 5-OxoETE, and (d5) resolvin D3. Samples and vial rinses (5% MeOH; 2 mL) were applied to Strata-X SPE columns (Phenomenex), previously washed with methanol (2 mL) and then dH_2_O (2 mL). Eicosanoids eluted with isopropanol (2 mL), were dried *in vacuuo* and reconstituted in EtOH:dH_2_O (50:50;100 μL) prior to ultra-high performance liquid chromatography electrospray ionization-MS/MS (UPLC ESI-MS/MS) analysis.

The eicosanoids were separated using a Shimadzu Nexera X2 LC-30AD coupled to a SIL-30AC auto injector, coupled to a DGU-20A5R degassing unit in the following way. A 14 min, reversed phase LC method utilizing an Acentis Express C18 column (150 mm x 2.1 mm, 2.7 µm) was used to separate the eicosanoids at a 0.5 mL/min flow rate at 40°C. The column was equilibrated with 100% Solvent A [acetonitrile:water:formic acid (20:80:0.02, v/v/v)] for 5 min and then 10 µL of sample was injected. 100% Solvent A was used for the first two min of elution. Solvent B [acetonitrile:isopropanol:formic acid (20:80:0.02, v/v/v)] was increased in a linear gradient to 25% solvent B at 3 min, to 30% at 6 min, to 55% at 6.1 min, to 70% at 10 min, to 100% at 10.10 min, 100% solvent B was held constant until 13.0 min, where it was decreased to 0% solvent B and 100% Solvent A from 13.0 min to 13.1 min. From 13.1 min to 14.0 min. Solvent A was held constant at 100%. The eicosanoids were analyzed via mass spec using an AB Sciex Triple Quad 5500 Mass Spectrometer. Q1 and Q3 were set to detect distinctive precursor and product ion pairs. Ions were fragmented in Q2 using N2 gas for collisionally-induced dissociation. Analysis used multiple-reaction monitoring in negative-ion mode. Eicosanoids were monitored using precursor → product MRM pairs. The mass spectrometer parameters used were: Curtain Gas: 20 psi; CAD: Medium; Ion Spray Voltage: −4500 V; Temperature: 300 °C; Gas 1: 40 psi; Gas 2: 60 psi; Declustering Potential, Collision Energy, and Cell Exit Potential vary per transition.

#### TUNEL analyses

Human islets (250/condition) were treated with cytokines or PGs, as described above, for 24h and then processed for TUNEL analyses

#### ChIP Analyses

Human islets (3500-5000/condition) were treated with cytokines for 2h or PGs for 4h, as described above, and processed for ChIP using antibodies select for p65 or iPLA_2_β and subsequent qPCR analyses for *PLA2G6*, *p65-NF*κ*B*, and *GcG* promoter regions.

### Statistical Analysis

All statistical analyses were done using GraphPad Prism (version 10). A “p” value < 0.05 was defined as significant. Student’s t-test was used for comparisons between 2 groups. For lipidomics analyses, “p” values were determined by two-way ANOVA, followed by Dunnett’s multiple comparisons test.

## Results

### iPLA_2_β induction is mitigated by inhibition of cytokine-induced ER Stress

β-cell ER stress precedes T1D development and inflammatory cytokines play critical roles in this process. We reported that ER stress induces iPLA_2_β expression increased in ER-stressed β-cells^8^ and that targeting iPLA_2_β reduces T1D incidence^11^. In the present study, we sought to examine the molecular mechanisms that lead it increases in iPLA_2_β expression. To address this, we first examined whether mitigation of ER stress impacts iPLA_2_β expression. We find that proinflammatory cytokines-induced INS-1 β-cell apoptosis is prominent after 24h (**Fig. 1A**) and that this was associated with induction of iPLA_2_β and ER stress factors (pPERK and CHOP) (**Figs. 1B-E**, cumulative densitometries are presented in **Suppl. Fig. 1**). As expected, the chemical chaperon PBA reduced induction of pPERK and CHOP, and β-cell apoptosis. Moreover, it reduced iPLA_2_β expression, suggesting a causative link between ER stress and iPLA_2_β expression in β-cells. The participation of NFκB in β-cell death leading to T1D is well established and we find a time-dependent nuclear accumulation of p65-NFκB following exposure to cytokines that peaked at 16h (**Fig. 1F-G**). However, in the presence of PBA, there was reduction in p65-NFκB (**Fig. 1H**). Given the established roles of both NFκB and STAT1 as transcriptional inducers in β-cells during T1D development, we considered the possibility that their activity in stressed β-cells is linked to iPLA_2_β expression.

### ER-stressed-mediated activation of NFκB and STAT1 in β-cells is linked to iPLA_2_β expression

Given the transcriptional activity of p65-NFκB and STAT1, we assessed whether ER-stressed induces iPLA_2_β through an NFκB-and/or STAT1-dependent mechanism. β-cells were pre-treated with Bay11-7082 or AZD 1480 (inhibitors of NFκB and STAT1 activation, respectively) prior to addition of cytokines. We find that both Bay11 (**Figs. 2A,B-F**) and AZD (**Figs. 2G, H-I**) reduced p65-NFκB and pSTAT1 expression, as well as pPERK and CHOP (cumulative densitometries are presented in **Suppl. Fig. 1**) and these outcomes were associated with reduced β-cell apoptosis (**Fig. 1J**). Further, cytokine-induced iPLA_2_β expression was also reduced in the presence of the two inhibitors. These findings suggest that iPLA_2_β induction in ER-stressed β-cells occurs via NFκB and STAT1-dependent pathways.

**Figure 2.**
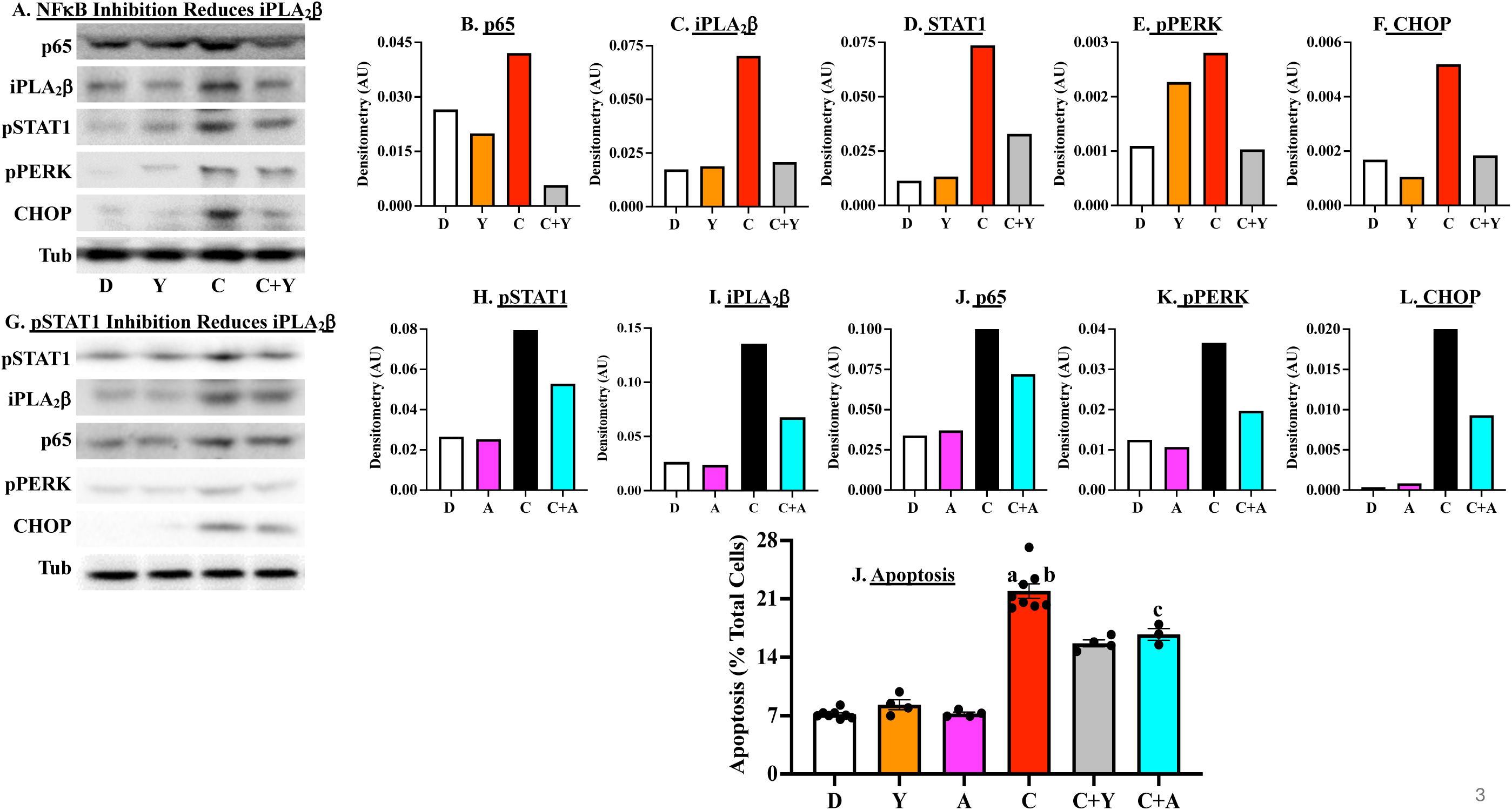
Inhibition of NFκB and STAT1 Reduces ER Stress-Mediated iPLA_2_β Induction and β-Cell Apoptosis. **A**. MIN-6 cells were pre-treated with NFκB inhibitor Bay11-7082 (10 µM) or STAT1 inhibitor AZD 1480 (10 µM) for 1h. The cells were then treated with <u>D</u>MSO or <u>c</u>ytokines (50 U/ml IL-1β + 150 U/ml IFNγ) for 16h and processed for p65-NFκB, SpTAT1, iPLA_2_β, pPERK, CHOP, and loading control tubulin immunoblotting analyses. Representative blots after BAY11 treatment (**A**) and corresponding densitometries (**B-F)**). Representative blots after AZD treatment (**G**) and corresponding densitometries (**H-I**). Cumulative densitometries are presented in **Suppl. Fig. 1. J**. Cells treated for 24h were processed for TUNEL analyses. (^a^Significantly different from DMSO, p < 0.0001; ^b^Significantly different from CTK+Bay11, p = 0.0007; ^c^Significantly different from CTK+AZD, p = 0.0028.)

In their roles as transcription factors, p65-NFκB and pSTAT1 induce genes associated with inflammation and apoptosis. We therefore considered the possibility that iPLA_2_β expression during ER stress is through p65-NFκB and/or pSTAT1 at the transcriptional level. Survey of the *Pla2g6* gene identified putative promoter regions containing binding motifs that are recognized by p65-NFκB and pSTAT1. This raised the possibility that p65-NFκB and pSTAT1 can induce transcription of *Pla2g6*. To address this, INS-1 cells were treated with cytokines and then processed for CHIP protocol using 1° antibodies targeting p65, STAT1, acetylated histone (AcH4), or IgG. The IPed DNA was then processed for qPCR analyses for *IL-1* (positive control) and *Pla2g6* promoter regions. Preliminary studies suggested peak effects with 2h cytokine treatment (**Suppl. Fig. 2**). We find exposure to cytokines promotes enrichment of p65-NFκB and pSTAT1 at the *IL-6* promoter region in β-cells (**Fig. 3A**) confirming triggering of transcriptional activation by cytokines and verified by AcH4 enrichment at the *IL-6* promoter region. Under the same conditions, p65-NFκB and pSTAT1, and AcH4 were also found to be enriched at the *Pla2g6* promoter region (**Fig. 3B**). Specificity of p65-NFκB, pSTAT1, and AcH4 enrichment at both *IL-6* and *Pla2g6* promoter regions was reflected by the absence of IgG (negative control) enrichment at the two promoter regions. These findings suggest that iPLA_2_β expression in ER-stressed β-cells is modulated of p65-NFκB and STAT1, in part, at the transcriptional level.

**Figure 3.**
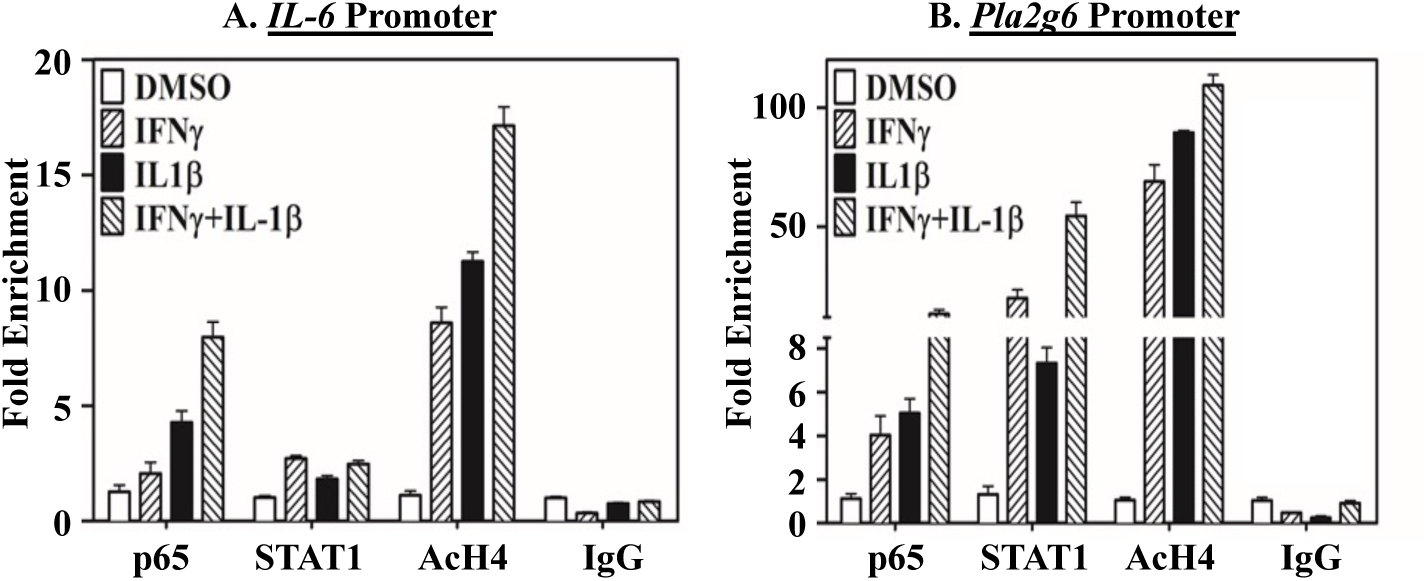
Transcriptional Induction of *Pla2g6* by NFκB and STAT1. INS-1 Cells were treated with 50 U/ml IL-1β + 150 U/ml IFNγ for 2h and processed for ChIP analyses. Nuclear preparations were immunoprecipitated with antibodies directed at p65, STAT1, AcH4, or IgG control and probed for association with *IL-6* (**A**) and *Pla2g6* (**B**) promoter regions by qPCR.

### Impact of iPLA_2_β on NFκB and STAT1 Activation in ER Stressed β-Cells

As targeting iPLA_2_β reduced T1D incidence, we considered the impact of iPLA_2_β itself on ER stress at the molecular level. To address this, we compared the effects of selective inhibition of iPLA_2_β, iPLA_2_γ, and cPLA_2_α. We find that inhibition of iPLA_2_β with either *S*-BEL^55^ (**Fig. 4A**) or FKGK18 (**Fig. 4B**)^56,57^ was effective in reducing p65-NFκB, pSTAT1, pPERK, and CHOP. However, inhibition of iPLA_2_γ with *R*-BEL^55^ (**Fig. 4C**) or cPLA_2_α with Cay10502 (**Fig. 4D**) did not lead to reductions in p65-NFκB, pSTAT1, pPERK, or CHOP. (Cumulative densitometries are presented in **Suppl. Fig. 3**). These findings suggest a select impact of iPLA_2_β in ER-stress β-cells.

**Figure 4.**
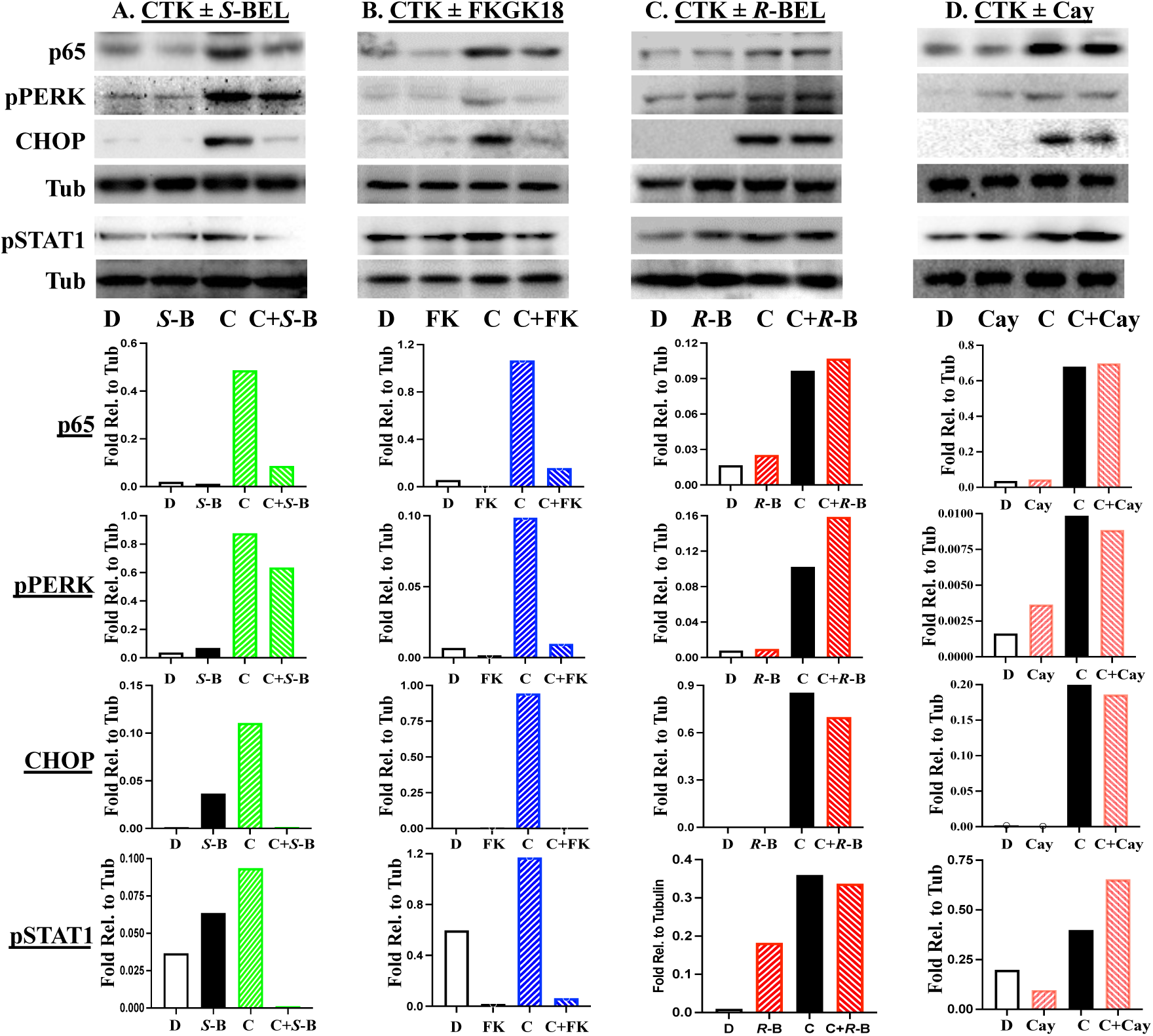
Selective Impact of iPLA_2_β Inhibition on NFκB and STAT1 Activation in ER Stressed β-Cells. MIN6 β-cells were pre-treated for 1h with selective inhibitors for iPLA_2_β (**A**) *<u>S</u>*<u>-B</u>EL (1 µM) or (**B**) <u>FK</u>GK18 (5 x 10^-8^ M); iPLA_2_γ (**C**) *<u>R</u>*<u>-B</u>EL (1 µM); or cPLA_2_α (**D**) Cay10502, (10 µM) and then treated for 16h with DMSO, or <u>C</u>ytokines (50 U/ml IL-1β + 150 U/ml IFNγ). The cells were then processed for p65-NFκB, pPERK, CHOP, pSTAT1, and loading control tubulin immunoblotting. Representative blots and densitometries are presented. Cumulative densitometries are presented in **Suppl. Fig. 3**.

To verify that such an impact is specifically iPLA_2_β-dependent, siRNA was used to knock down *Pla2g6* mRNA and the cells were treated with vehicle or cytokines. While the cytokines induced iPLA_2_β, p65, pSTAT1, pPERK, and CHOP in cells transfected with scrambled RNA (scrRNA), they were not in cells transfected with siRNA targeting *Pla2g6* (**Fig. 5**). Further, p65-NFκB induction evident with scrRNA was also reduced with siRNA treatment (**Figs. 5A,C**), in the absence of reductions in IκBα (**Figs. 5D**). (Cumulative densitometries are presented in **Suppl. Fig. 3**). Consistently, only inhibition of iPLA_2_β (with *S*-BEL or FKGK18) or iPLA_2_β knockdown (with siRNA), but not inhibition of iPLA_2_γ or cPLA_2_α, mitigated cytokine-induced β-cell apoptosis (**Fig. 6**). These findings suggest a select impact of iPLA_2_β on ER stress outcomes in β-cells, including activation of NFκB and STAT1.

**Figure 5.**
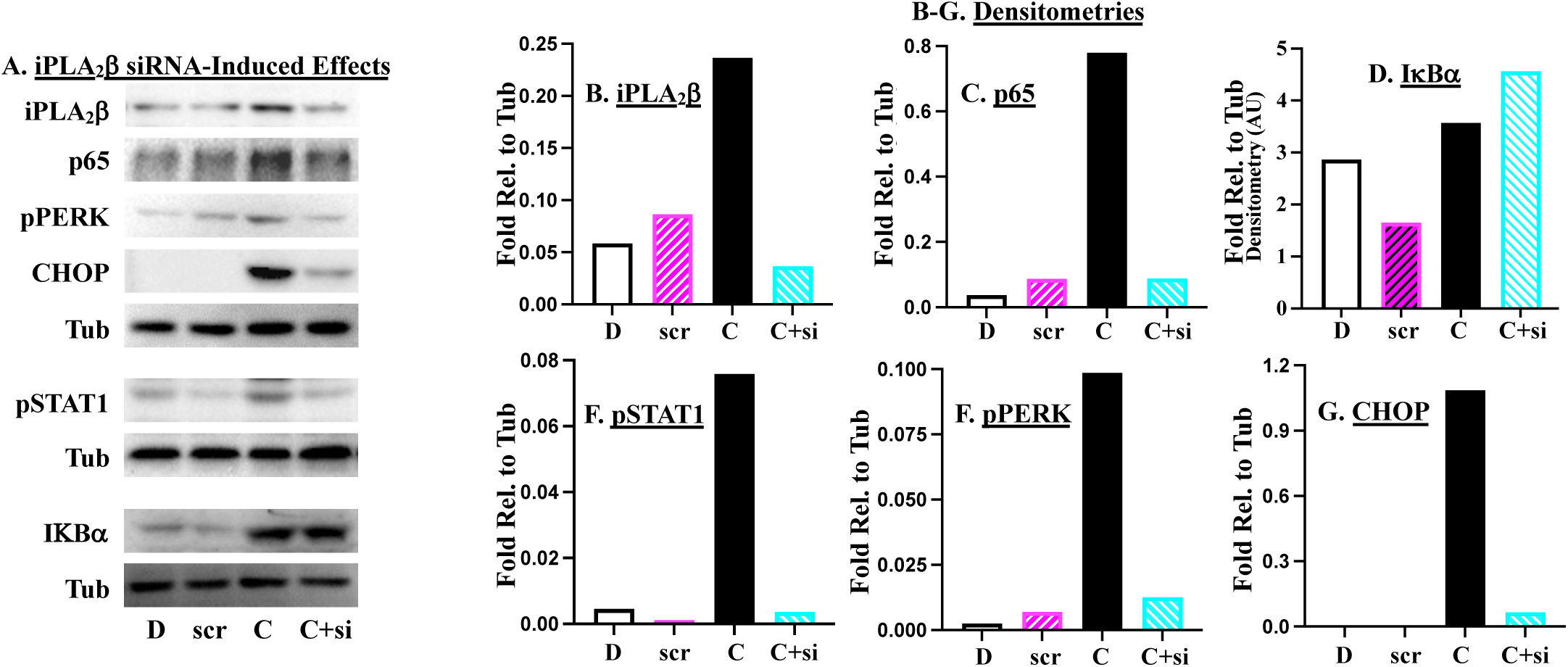
Knockdown of iPLA_2_β Reduces ER Stress and NFκB and STAT1 Activation in ER Stressed β-Cells. MIN6 β-cells were transfected with <u>scr</u>ambled RNA or <u>si</u>RNA targeting *Pla2g6* and treated with <u>D</u>MSO or <u>C</u>ytokines (50 U/ml IL-1β + 150 U/ml IFNγ) and processed for iPLA_2_β, p65-NFκB, IKBα, pSTAT1, pPERK, CHOP, and loading control tubulin immunoblotting. Representative blots (**A**) and corresponding densitometries (**B-G**) are presented. Cumulative densitometries are presented in **Suppl. Fig. 3**.

**Figure 6.**
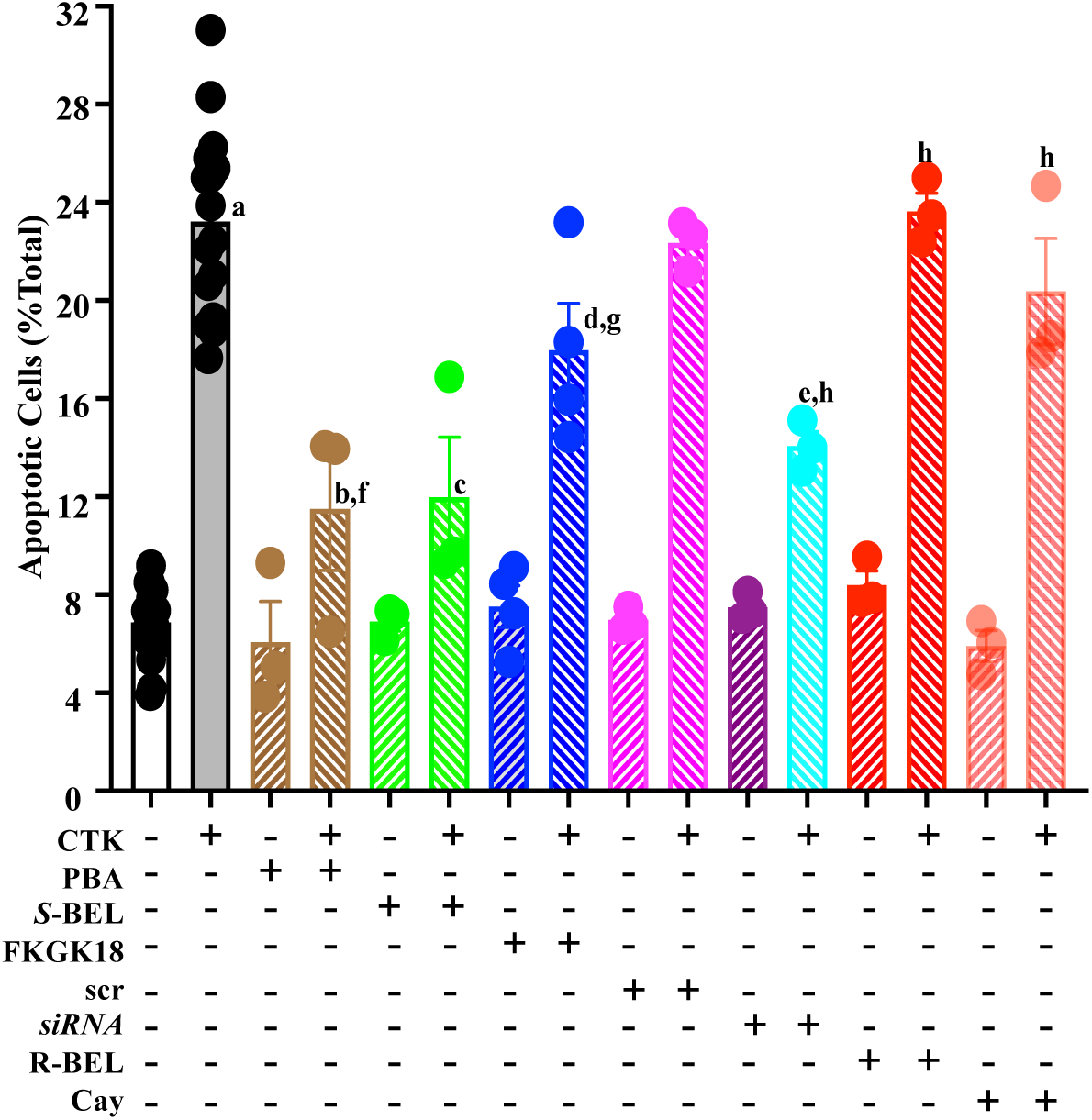
Select Targeting of iPLA_2_β Inhibits ER Stress-Induced β-Cell Apoptosis. **A.** MIN6 β-cells were pretreated with *<u>S</u>*<u>-B</u>EL (1 µM), <u>FK</u>GK18 (5 x 10^-8^ M), *<u>R</u>*<u>-B</u>EL (1 µM), or Cay10502 (10 µM) for 1h or transfected with <u>scr</u>ambled or siRNA targeting *Pla2g6* and then treated with <u>D</u>MSO or <u>c</u>ytokines (50 U/ml IL-1β + 150 U/ml IFNγ) for 24h and processed for TUNEL analyses. Data are means±SEMs of % apoptotic cells, relative to total cells. (^a^Cytokine group significantly different from DMSO group, p<0.0001; ^b,c,d,e,f,g,h^Cytokine group significantly different from cytokines + inhibitor or siRNA groups: p=0.0001, p=0.0002, p=0.0228, p=0.0007, p=0.0010, p=0.0004, p=0.0001, respectively.

### CRISPR-Cas9 knockdown of iPLA_2_β reduces its association with promoters

We previously reported that during ER stress, iPLA_2_β translocates to the nucleus^8^ and in view of the above findings, we considered the possibility iPLA_2_β modulates NFκB at the gene level. To address this question, we established a CRISPR-Cas9 approach to knockdown iPLA_2_β. Using Benchling ^(^www.benchling.com), we designed two CRISPR guides (G1 and G2) targeting the coding region in exon 2 (**Fig. 7A**). Following the method described by Ran et al^58^, we generated plasmid constructs that express the guide RNAs along with Cas9 protein fused either to EGFP or puromycin to facilitate screening and selection. We then transfected MIN6 β-cells with either control plasmids (G0) or plasmids encoding gRNAs (G1 and G2) targeting *Pla2g6*; in addition to indels at each of the two target sites, simultaneous cutting of both target sites followed by NHEJ repair would result in a predicted deletion. We performed genotyping by PCR followed by TBE-PAGE analyses to confirm the presence of the deletion allele (**Fig. 7B**). Whereas only the expected wildtype band (298 bp) is predominantly expressed in G0-treated (control) cells, the cells transfected with both the CRISPR plasmids (G1 and G2) had additional truncated bands, in particular one at 161bp, which is expected from the 137 bp deletion by synchronous activity of both CRISPR guides. Additional minor bands that are uniquely found in the CRISPR targeted samples are likely to be alleles with different deletions arising from NHEJ, which is a relatively common phenomenon^59^. The 137 bp deletion leads to a frameshift in the coding region resulting in an early truncation. The mutant allele encodes a 30 aa peptide with only 8 native amino acids at the N-terminal and 22 new amino acids resulting from the frameshift (**Fig. 7C**). Immunoblotting analyses verified the knockdown of basal and cytokine-induced iPLA_2_β with the CRISPR constructs (G1 and G2), but not with the control plasmids (G0) (**Fig. 7D,E**). Consistently, cytokine-induced β-cell apoptosis was unaffected by G0 but reduced with the CRISPR targeting (**Fig. 7F**). As both gRNAs appeared to work similarly, they were used in combination in subsequent experiments. Such studies demonstrated reductions in cytokine-induced iPLA_2_β, p65-NFκB, pSTAT1, and pPERK with CRISPR targeting, relative to controls (**Figs. 7G-I**). These findings provide evidence of successful application of the CRISPR-Cas9 protocol to knockdown iPLA_2_β in β-cells, resulting in the mitigation of ER stress outcomes.

**Figure 7.**
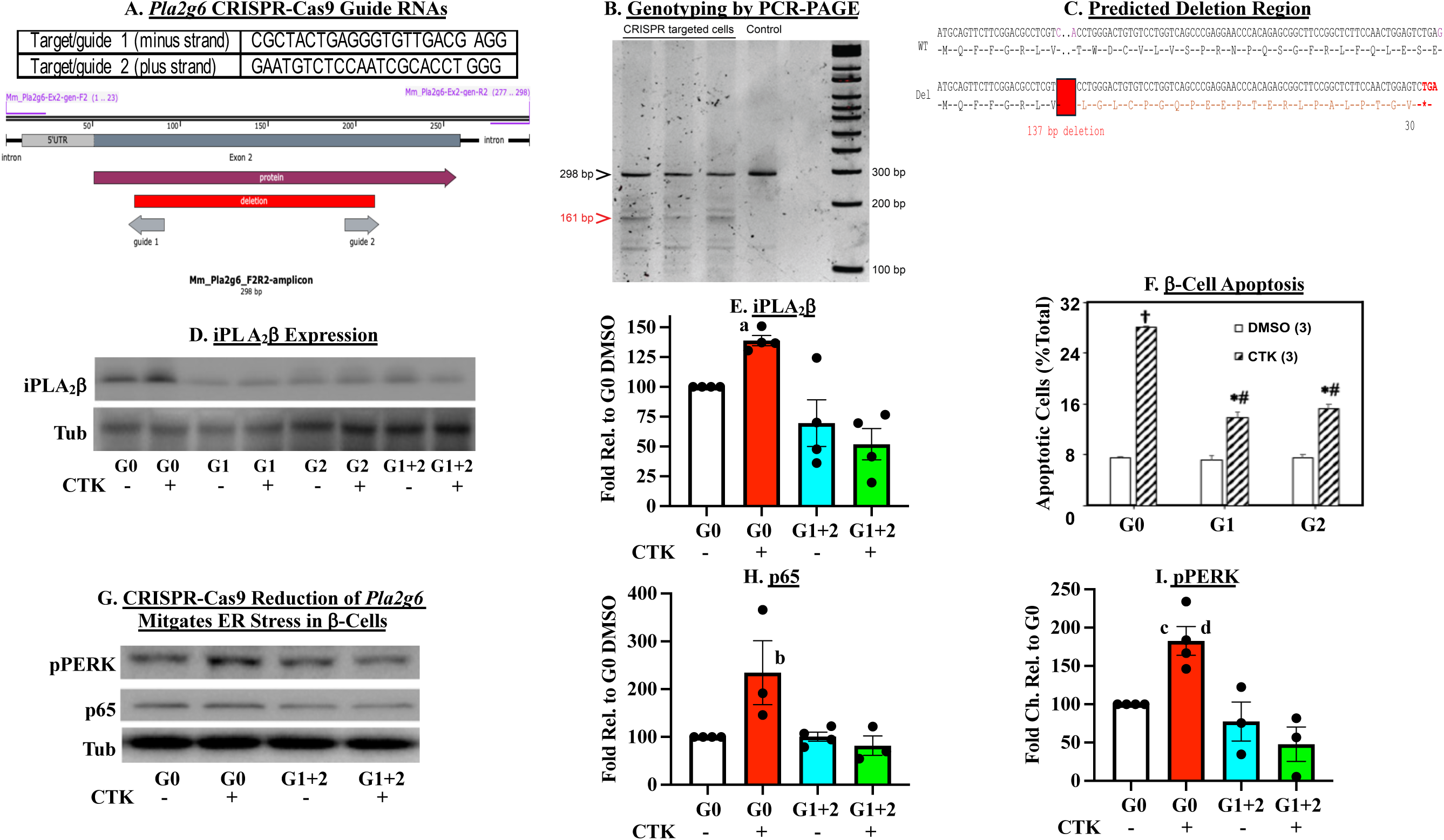
CRISPR-Cas9 Approach to Examine iPLA_2_β Role in Transcription. MIN6 β-cells were transfected with control (G0) or guide RNA constructs (G1 and G2) targeting *Pla2g6*. **A.** *Pla2g6* CRIPR-Cas9 gRNAs. **B**. Genotyping analyses by PCR. **C**. Predicted deletion region. **D, E**. Cells were treated with <u>D</u>MSO or <u>C</u>ytokines (50 U/ml IL-1β + 150 U/ml IFNγ) for 16h and then processed for iPLA_2_β and loading control tubulin immunoblotting. Representative blots and densitometries are presented. **F**. Cells were treated with <u>D</u>MSO or <u>C</u>ytokines (50 U/ml IL-1β + 150 U/ml IFNγ) for 24h and then processed for TUNEL analyses. Data are means±SEMs of percent apoptotic cells, relative to total cells. (^†,*^Significantly different from G0-DMSO, p<0.00001, p<0.005; ^#^different from G0-CTK p<0.0005, n=3/group). **G-I**. Cells were treated with <u>D</u>MSO or <u>C</u>ytokines (50 U/ml IL-1β + 150 U/ml IFNγ) for (16h) and then processed for pPERK, p65-NFκB, and loading control tubulin immunoblotting and representative blots (**G,** n=3-4) and cumulative densitometries presented (**H-I**). ^a,b,c^G0 CTK group significantly from GO DMSO group, p<0.0001, p=0.05, p=0.0046, respectively; ^d^G0 CTK group significantly different from G1+G2 CTK group, p<0.0057.)

### Evidence for iPLA_2_β impact at the transcription level

To determine if iPLA_2_β modulates NFκB or STAT1 at the transcription level, CRISPR-Cas9-modified MIN6 cells were treated with DMSO vehicle or cytokines for 2h, as determined in pilot studies (**Suppl. Fig. 2**). After 2h, a ChIP protocol was performed using antibodies directed against iPLA_2_β, AcH4, and IgG. The IPed DNA was then used for qPCR analyses of *Pla2g6* and *Nf*κ*b* promoter regions. Intriguingly, we find enrichment of iPLA_2_β at both *Pla2g6 and Nf*κ*b* promoter regions under both basal and cytokine-exposed conditions in both the G0 and gRNAs-treated, relative to IgG enrichment (**Figs. 8A,B**). Under the same conditions, enrichment of AcH4 at the *Pla2g6* promoter region was also evident, indicative of triggering of *Pla2g6* transcription. Interestingly, enrichment of iPLA_2_β at the *Pla2g6* and *Nf*κ*b* promoter regions or of Ach4 at the *Pla2g6* promoter region were enhanced by cytokines, relative to DMSO, in the G0 group but not in the gRNA-treated group (**Figs. 8C**). These findings suggest a correlation between iPLA_2_β abundance and its association with promoters and ability to modulate transcriptional activity.

**Figure 8.**
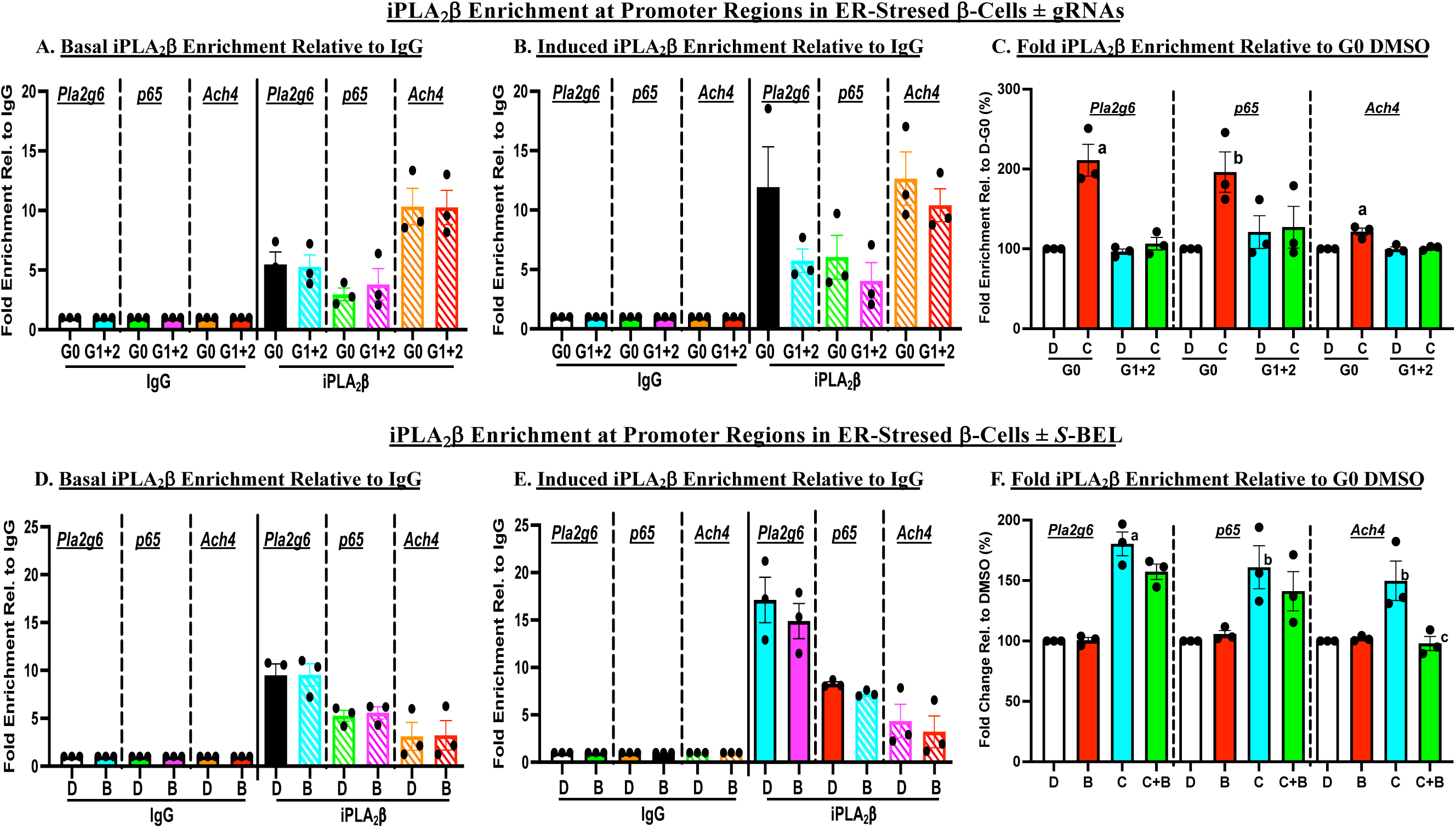
iPLA_2_β Associates with *Pla2g6* and *p65* Promoters in ER Stressed β-Cells. MIN6 β-cells were transfected with control (G0) or guide RNA constructs (G1 and G2) targeting *Pla2g6* (**Top panels**) or were pretreated with *S*-<u>B</u>EL (10 µM, 1h) (**Bottom panels**) and then treated with <u>D</u>MSO or <u>C</u>ytokines (50 U/ml IL-1β + 150 U/ml IFNγ) for (2h). Subsequently, ChIP was performed using antibody directed against iPLA_2_β and acetylated histone (AcH4) followed by qPCR analyses of *Pla2g6* and *Nf*κ*b* promoter regions. AcH4 was used to confirm transcriptional activation and IgG as a negative control. **A, D.** Fold enrichment, relative to IgG, under basal (DMSO) conditions. **B, E.** Cytokine-induced fold enrichment, relative to IgG. **C, F.** Cytokine-induced fold enrichment, relative to DMSO: **C**. **^a,b^**G0 C groups significantly different from other groups, p<0.01, p<0.05, respectively; n=3.); **F**. **^a,b^**Cytokine groups significantly different from corresponding DMSO groups, p<0.005, p<0.05, respectively; **^c^**C+B significantly different from corresponding C p<0.05; n=3.)

Because iPLA_2_β is known to not contain any DNA binding motifs, the above findings were unexpected. However, as iDL signaling participates in ER stress-induced β-cell death, we anticipated that inhibition of iPLA_2_β may impair its association with promoters. We therefore performed ChIP analyses as above, but this time compared the MIN6 cells response in the absence and presence of *S*-BEL. As above, we find enrichment of iPLA_2_β at the *Pla2g6*, *Nf*κ*b*, and of AcH4 at the *Pla2g6* promoter regions under both basal (**Fig. 8D**) and cytokine-exposed (**Fig. 8E**) conditions in the absence and presence of *S*-BEL, relative to IgG. Surprisingly, in contrast to the above findings in iPLA_2_β-reduced MIN6 cells, enrichment of iPLA_2_β at the *Pla2g6* and *Nf*κ*b* promoter regions was similar in the absence and presence of *S*-BEL (**Fig. 8F**). However, Ach4 enrichment at the *Pla2g6* promoter region was decreased, suggesting mitigation of transcriptional activity. These findings raise the possibility of a unique separation of association with a promoter region from activation of transcription, involving lipid signaling.

### ER stress-induced production of prostaglandins is reduced by inhibition of iPLA_2_β

We previously reported dramatic changes in proinflammatory eicosanoids profile in immune cells during T1D development^12^. A potential explanation for the above findings may be related to the abundances of IDLs in the β-cells, whose production would be limited by iPLA_2_β activation. To address this possibility, we performed lipidomics analyses to identify changes in lipid abundances associated with ER stress. As insulinoma cells generated weak lipid intensity signals and the required temporal analyses necessitated large batches of islets, we utilized human islets for these analyses. We previously reported that cytokines induce ER stress and human β-cells apoptosis that are mitigated by *S*-BEL^5^. Here, we report that under the same conditions, islets production of several proinflammatory eicosanoids increases in a temporal fashion that peaks at 24h. Among those, the production of PGE_2_, PGF_2_α, and 8-Iso-PGF_2_α is blunted by the addition of *S*-BEL (**Figs. 9A-C**). These PGs are noted for their potent pro-inflammatory effects^38^ raising the possibility of that their signaling contributes to ER stress-induced β-cell apoptosis.

**Figure 9.**
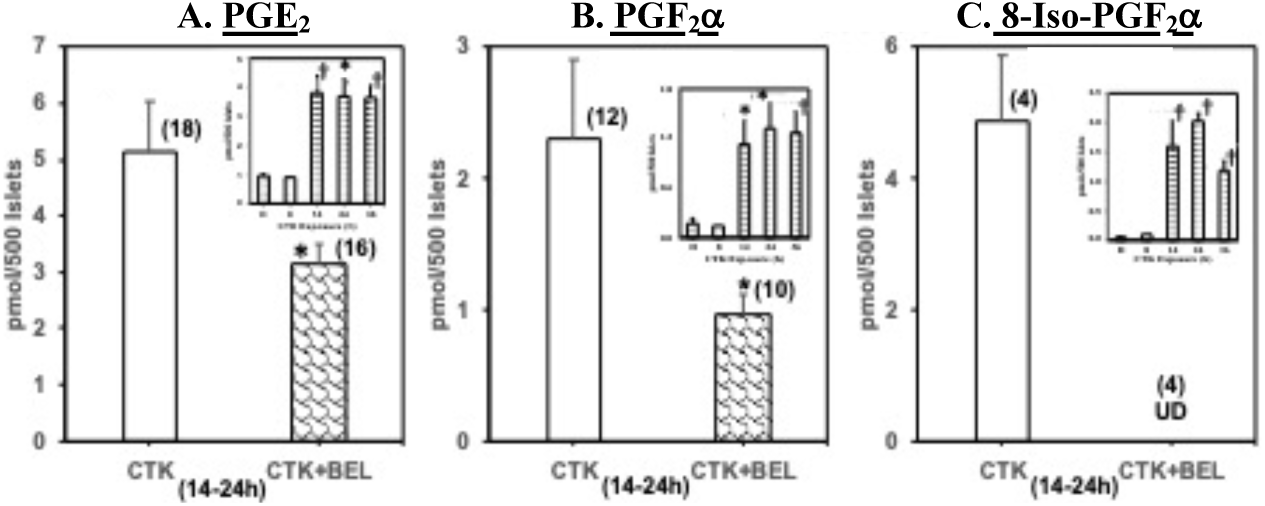
Lipidomics analyses. Human islets (500/condition) were treated with cytokines (100 U/ml IL-1β + 300 U/ml IFNγ) for 2-36h. In some replicates, at peak response times, cytokines + *S*-BEL groups were added. A. PGE_2_. B. PGF_2_α. C. 8-Iso-PGF_2_α. (Insets,^*,†^significantly different from DMSO group, p < 0.05 and p < 0.01, respectively, n=10-18. *CTK+*S*-BEL group significantly different from CTK group, p < 0.05, n values indicated above the bars.)

### Select prostaglandins contribute to ER stress and apoptosis

To address this possibility, MIN6 cells were treated with PGE_2_, PGF_2_α, and 8-Iso-PGF_2_α alone or in combination for 24h and then processed for TUNEL analyses. We find that each PG alone induced MIN6 β-cell apoptosis, relative to DMSO (**Supplemental Fig. 4**) that was ∼50% of that caused by cytokines. And, in combination they induced ∼67% of that caused by cytokines (**Fig. 10A**). Further, the combination of the three PGs induced p65-NFκB, pSTAT1, pPERK, and CHOP, relative to DMSO-treated cells (**Fig. 10B-F**). These findings suggest that PGE_2_, PGF_2_α, and 8-Iso-PGF_2_α signaling participates in ER stress-mediated β-cell apoptosis.

**Figure 10.**
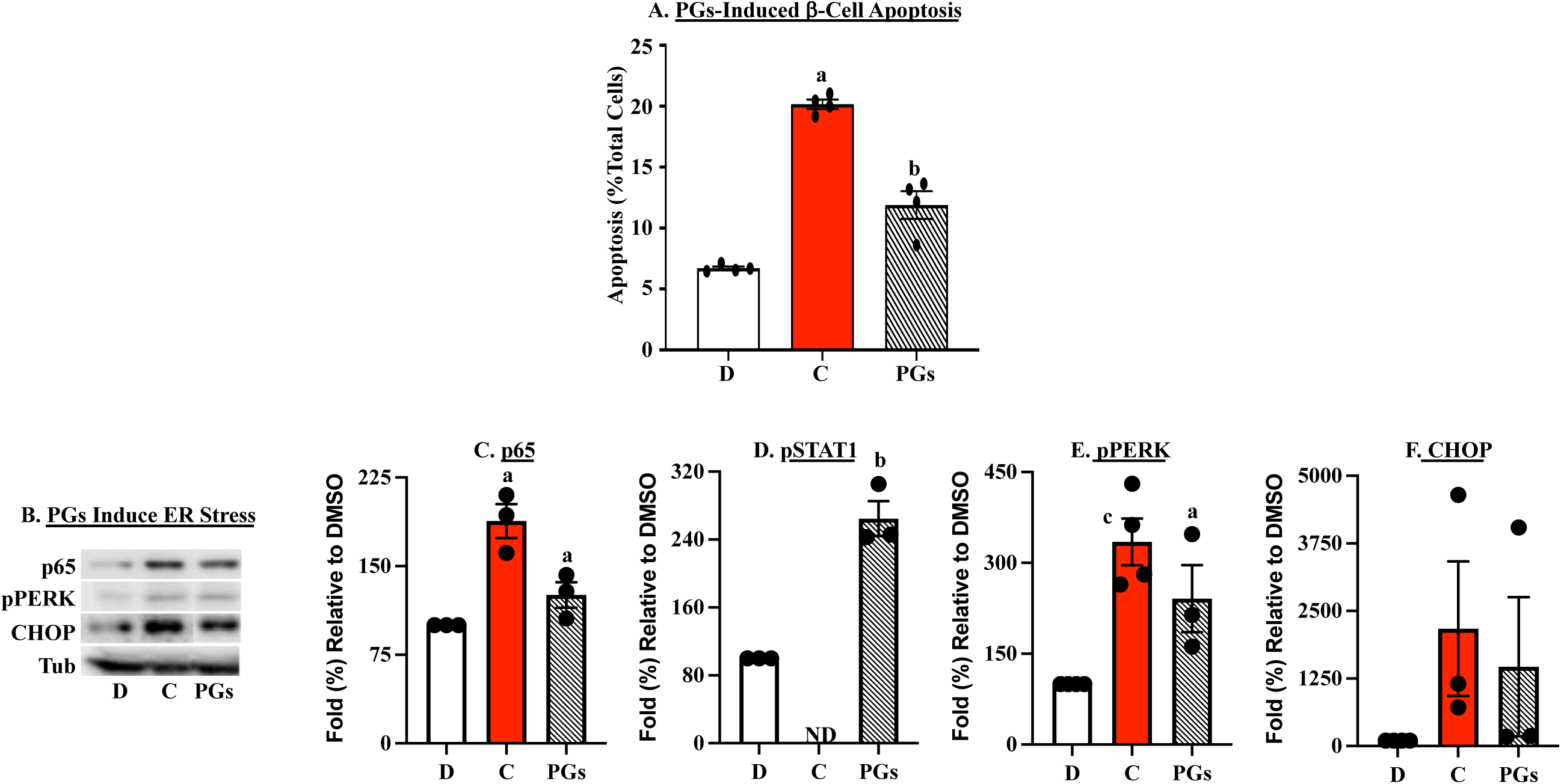
Impact of Select PGs β-Cell Apoptosis and ER Stress. **A.** MIN6 β-cells were treated with <u>D</u>MSO, <u>C</u>ytokines as a positive control (50 U/ml IL-1β + 150 U/ml IFNγ) or a PGs cocktail (10 nM PGE_2_ + 10 µM PGF_2_α + 10 µM 8-Iso-PGF_2_α) and processed for TUNEL analyses after 24h. Data are means ± SEMs of % apoptotic cells, relative to total cells. (^a^Cytokine group significantly different from DMSO group, p < 0.0001; ^b^PGs group significantly different from DMSO group, p = 0.004, n=3/group). **B**. Representative immunoblotting analyses blots for p65-NFκB, pSTAT1, pPERK, CHOP, and loading control tubulin after 16h. **C-F**. Corresponding cumulative densitometries are presented. (^a,b,c^Cytokine or PGs group significantly different from DMSO groups, p, 0.05, p = 0.0013, and p = 0.009, respectively, n=3-4/group. ND=not determined.)

### Feedback regulation of iPLA_2_β expression

While regulation of enzymes by PGs has been reported^60^, their potential impact on β-cell iPLA_2_β has not. This possibility surfaced by the findings that while select inhibition of iPLA_2_β with *S*-BEL (**Fig. 11**) or FKGK18 (**Fig. 11B**) had a minor effect alone in MIN6 cells, they markedly reduced cytokine-induced iPLA_2_β expression. A select nature of iPLA_2_β effect was reflected by the absence of a similar reduction with inhibition of iPLA_2_γ or cPLA_2_α (**Fig. 11C,D**). Analogous to cytokine effects, the combination PG treatment also increased iPLA_2_β expression (**Fig. 11E**). (Cumulative densitometries are presented in **Suppl. Figs. 3 and 6**) These findings suggest that PGs production by stressed cells can induce iPLA_2_β expression.

**Figure 11.**
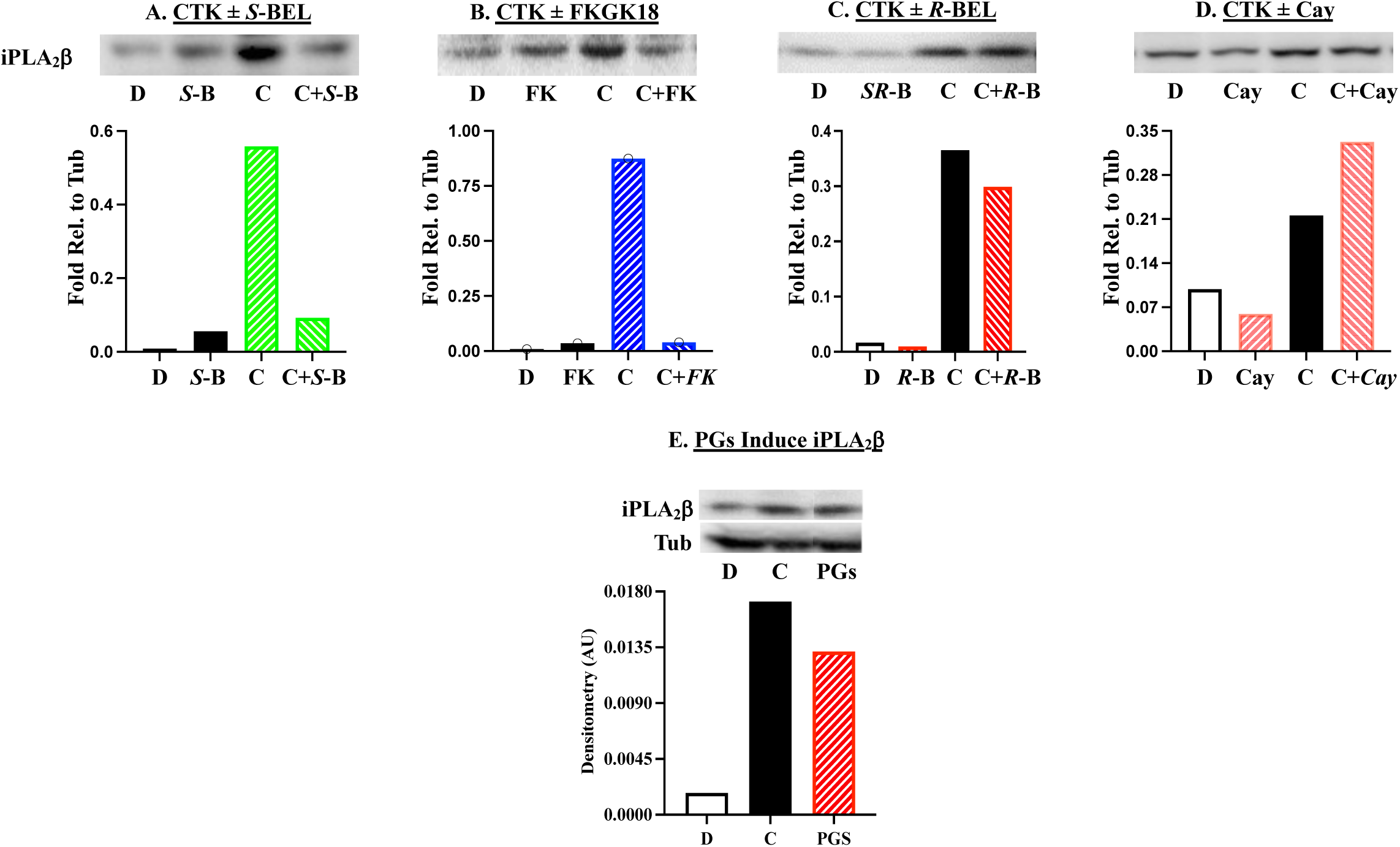
iDLs-Dependent Induction of iPLA_2_β. MIN6 β-cells were treated with <u>C</u>ytokines ± inhibitors as in Fig. 4, or PGs, as in Fig. 9, and processed for iPLA_2_β immunoblotting analyses. Representative blots and corresponding densitometries for CTK ± *S*-BEL (A), CTK ± FKGK18, CTK ± *R*-BEL, CTK ± *S*-BEL, and cytokines and PGs (E) are presented. Cumulate densitometries are presented in Supplemental Figures 3 and 6.

### Select prostaglandins promote iPLA_2_β association with promoters

Translocation of iPLA_2_β to the nucleus amplifies the activity of truncated iPLA_2_β resulting in accelerated hydrolysis of the *sn*-2 fatty acid from membrane glycerophospholipids and this would be expected to increase the abundances of iDLs in the nucleus. To assess whether the select PGs participate in transcriptional regulation to upregulate iPLA_2_β expression, we performed ChIP analyses with MIN6 cells treated with cytokines or the combination of PGE_2_, PGF_2_α, and 8-Iso-PGF_2_α in the absence or presence of *S*-BEL. Pilot studies indicated that 4h treatment produced peak effects (**Suppl. Fig. 7**) and subsequent analyses revealed that the combination PGs increased the association of iPLA_2_β with *Pla2g6*, *Nf*κ*b*, and *Stat1* promoter regions, relative to IgG (**Fig. 12A**). Further, the association of iPLA_2_β with *Pla2g6* (**Fig. 1B**), *Nf*κ*b* (**Fig. 1C**), and *Stat1* (**Fig. 1D**) following exposure to cytokines was significantly increased, relative to DMSO. Specificity of such enrichment was evidenced by the lack of PGs-induced enrichment of iPLA_2_β with *Gcg* (**Fig. 1E**) or *Pepck* (**Fig. 1F**) promoter regions. Addition of *S*-BEL did not have an effect alone or significantly impact stimulated association of iPLA_2_β with any promoter regions. These findings suggest a mechanism by which the PGs may confer upon iPLA_2_β the ability to activate transcription to not only promote gene product NFκB but, also iPLA_2_β. These findings suggest that PGs produced by stressed β-cells contribute to β-cell death by inducing iPLA_2_β, NFκB, and STAT1 expression at the transcription level.

**Figure 12.**
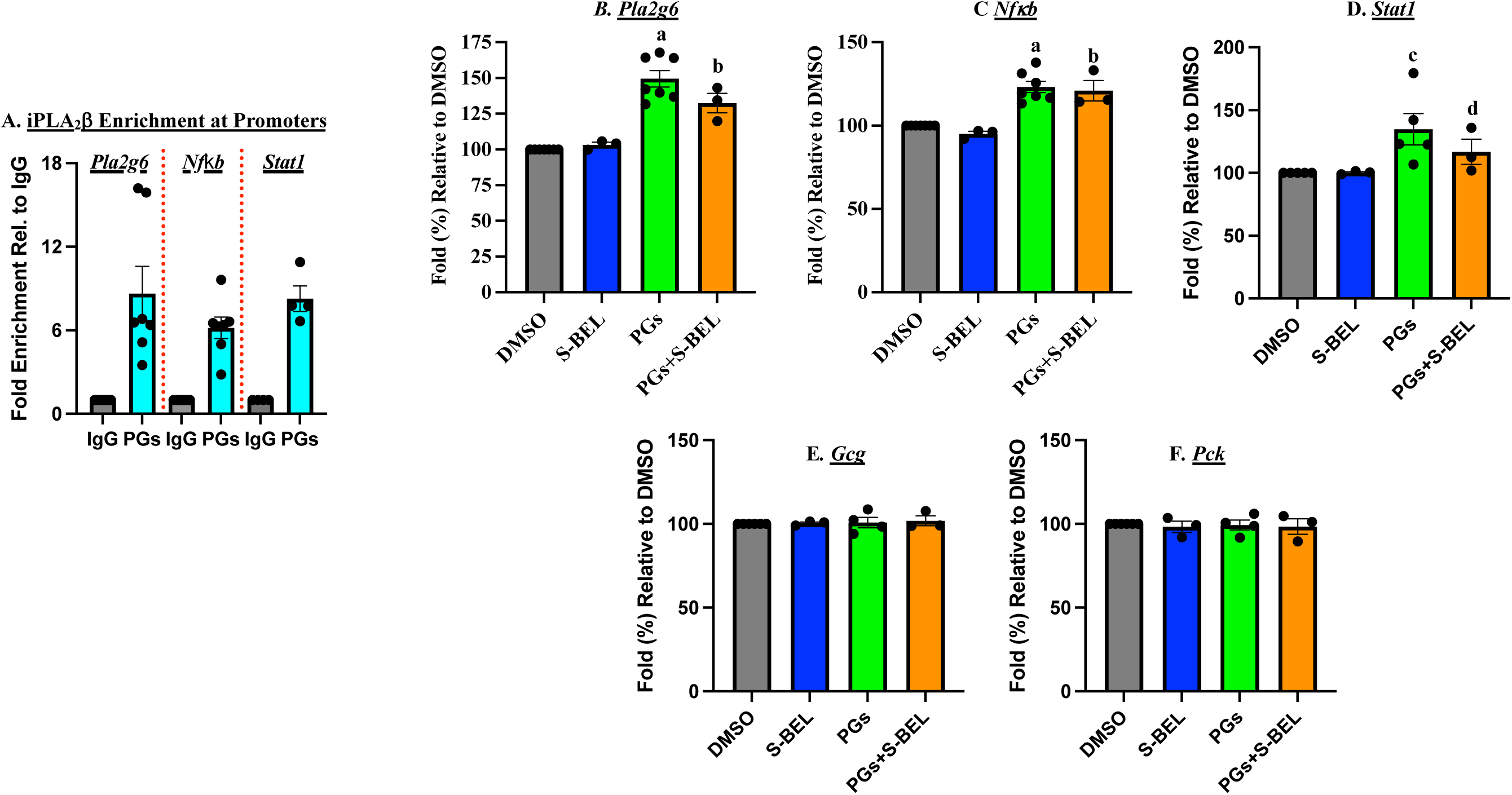
Select iDLs-Indued iPLA_2_β Enrichment at Promoter Regions in β-Cells. MIN6 β-cells were treated with a PGs cocktail, as in Fig. 10, for 4h and processed for ChIP and qPCR analyses. **A**. Fold enrichment of iPLA_2_β at promoter regions, relative to IgG. **B-F**. Fold enrichment of iPLA_2_β at promoter regions, relative to DMSO: **B**, *Pla2g6*; **C**, *NfkB*; **D**, *Stat1*; **E**. *Gcg*; **F**. *Pepck*. (^a,c^PGs groups significantly different from corresponding DMSO groups, p < 0.001 and p < 0.01, respectively. ^b,d^PGs + *S*-BEL groups significantly different from corresponding DMSO groups, p < 0.01 and p < 0.05, respectively.)

### Transcriptional regulation by select iDLs in human islets

To assess whether transcriptional regulation of and by iPLA_2_β translates to islet β-cells, ChIP analyses were performed in human islets. First, the association of p65 with *Pla2g6* promoter was confirmed by treating the islets with cytokines followed by a ChIP protocol using antibodies directed against p65-NFkB or IgG. Subsequent qPCR analyses revealed cytokine-induced enrichment of p65-NFκB at the *PLA2G6*, but not at the *GCG*, promoter region, relative to IgG (**Fig. 13A, Suppl. Fig. 7**). Also evident were cytokine-induced enrichment of iPLA_2_β at the *PLA2G6* and *NF*κ*B*, but not at the *GCG*, promoter regions, relative to IgG (**Fig. 13B**). Further, the select PGs also promoted enrichment of iPLA_2_β at the *PLA2G6* and *NF*κ*B*, but not at the *GCG*, promoter regions, relative to IgG (**Fig. 13C**). Consistently, the PGs combination induced a 2-fold increase in β-cell apoptosis, relative to DMSO (**Fig. 13D**). In comparison, the cytokines indued a 3-fold increase in β-cell apoptosis. These findings validate a link between iPLA_2_β/iDLs and transcriptional regulation of relevant apoptotic factors in human islet β-cells.

**Figure 13.**
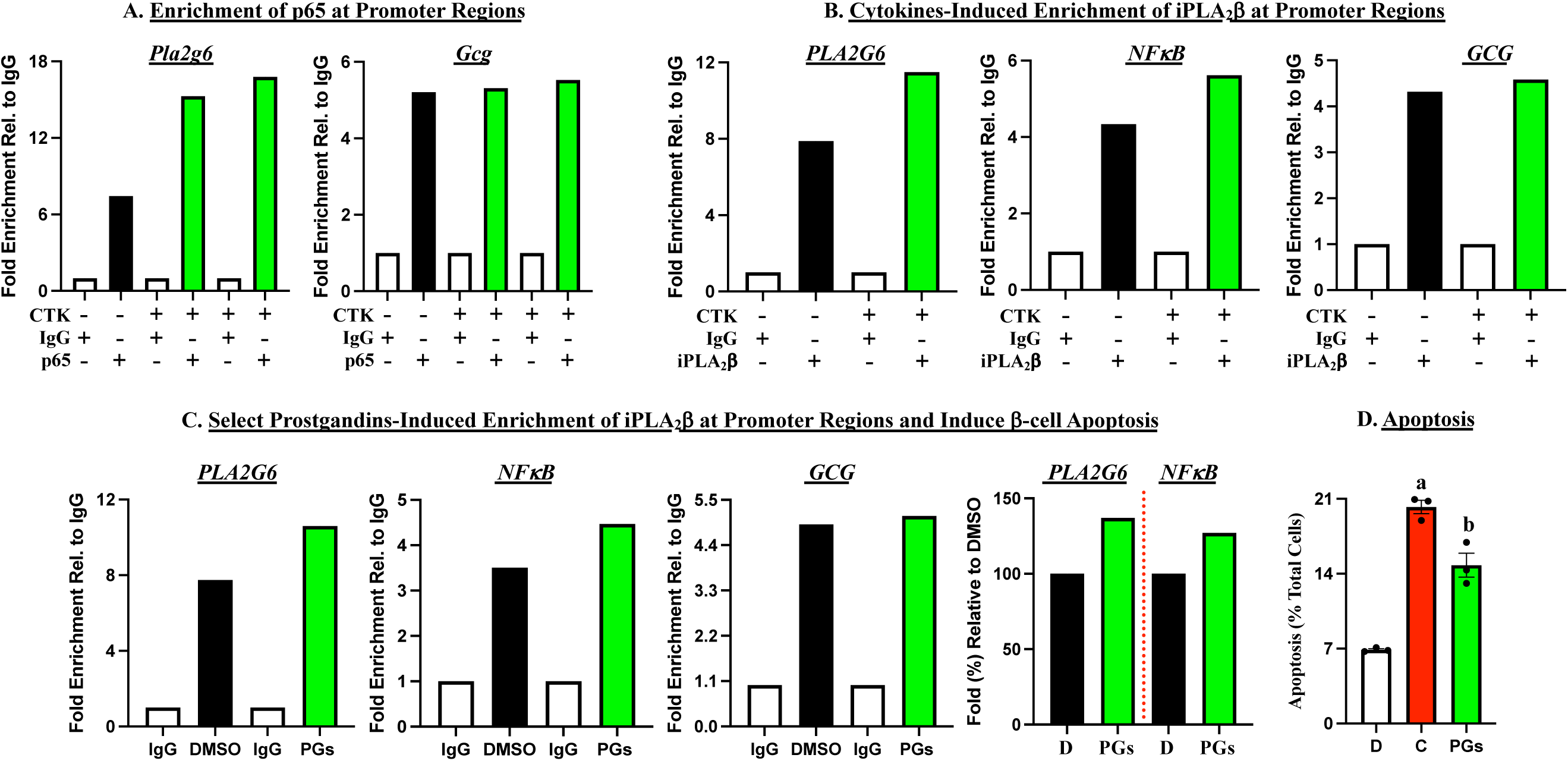
Validation of p65 or iPLA_2_β Enrichment at Promoter Regions in Human Islets. Human islets (3500-5000/condition) from healthy donors were treated with cytokines (100 U/ml IL-1β + 300 U/ml IFNγ for 2h (**A, B**) or a PGs cocktail (**C**) as in Fig. 10 for 4h. The islets were then processed for ChIP analyses using antibodies directed against p65-NFκB or iPLA_2_β and subsequent qPCR analyses. **D**. Human islets (250/condition) from healthy donors were treated with cytokines (100 U/ml IL-1β + 300 U/ml IFNγ or a PGs cocktail for 24h and processed for TUNEL analyses. Data are mean ± SEMs of %apoptotic cells, relative to total cells. (^a,b^Significantly different from DMSO group, p < 0.0001 and p = 0.004, respectively.)

**Figure 14.**
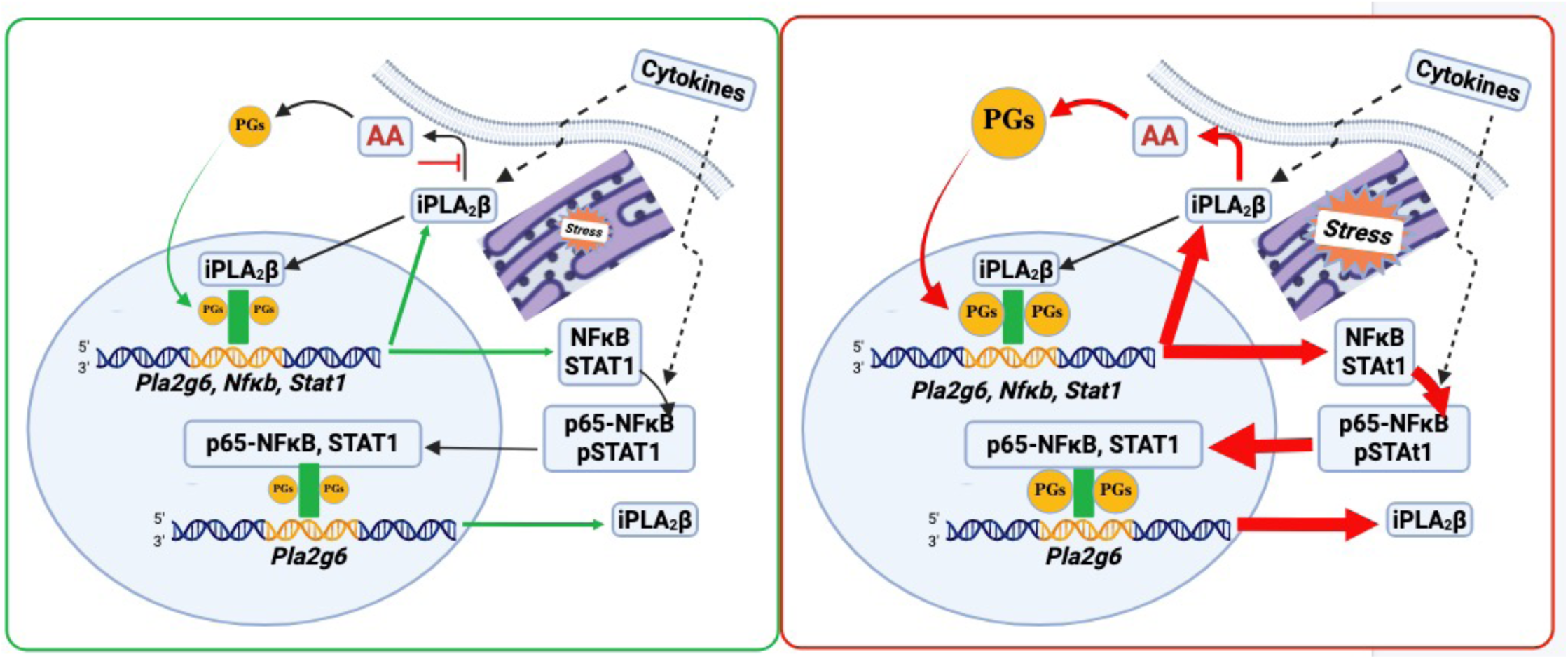
Proposed Model of iDLs Contribution to T1D Development.

## Discussion

T1D is a consequence β-cell destruction resulting from concerted actions of immune cells and β-cells. While the immunologic components have been heavily dissected, the role of β-cells in causing its own demise are not well-understood. We recently reported that select reduction of iPLA_2_β in macrophages or T-cells reduces their inflammatory phenotype and lowers T1D incidence^10,13^. However, the nature of signals derived from β-cells to drive this process are not completely understood.

Our work has implicated iPLA_2_β activation in β-cells as a critical contributor to ER stress-induced β-cell death^2–5,8^. Reports suggesting that ER stress in β-cells precedes T1D onset^29,61^ raises the possibility that iDL signaling is an early trigger leading to β-cell death and T1D. Activation of PLA_2_s leads to hydrolysis of the *sn*-2 substituent of membrane glycerophospholipids and islet β-cell membranes are enriched in *sn*-2 arachidonic acid^62^. Subsequent metabolism of arachidonic acid leads to generation of oxidized bioactive lipids including prostaglandins, several of which are potently inflammatory^38^. Here we sought to identify molecular mechanisms through which iDL signaling participates in ER stress-induced β-cell death. Utilizing insulinoma cells and human islets; employing chemical, siRNA, and CRISPR-Cas9 approaches; and immunoblotting, TUNEL, qPCR, ChIP, and lipidomics analyses, we report (a) a select role of iPLA_2_β in ER stress and β-cell apoptosis, (b) regulation of iPLA_2_β expression by NFκB and STAT1 at the transcriptional level, (c) regulation of NFκB, STAT1, and iPLA_2_β expression by iPLA_2_β at the transcriptional level, (d) potential co-factor roles of select inflammatory prostaglandins to facilitate transcriptional activation by iPLA_2_β, and (e) feedback regulation of iPLA_2_β expression.

ER stress elicits adaptive unfolded protein response (UPR) initially, however, prolonged ER stress can lead to terminal UPR, where hyperactivation of the UPR arms (PERK, IRE1α, ATF6) triggers apoptotic targets^63,64^. The nuclear transcription factors NFκB and STAT1, by virtue of being able to trigger pro-apoptotic genes expression are key players in cytokine-mediated β-cell death and T1D development^65–74^. Our finding that inhibiting NFκB or STAT1 activation decreases iPLA_2_β expression, in association with a decrease in β-cell apoptosis, led us to further explore the link between iPLA_2_β and the two transcription factors in ER stressed β-cells. Initial ChIP analyses with antibodies directed against NFκB and STAT1 revealed that cytokine exposure promotes association of NFκB and STAT1 at *Pla2g6* promoter region. While gene analyses identified evolutionarily conserved region in the *Pla2g6* promoter that harbors putative response elements for transcription factors that respond to ER stress, including NFκB and STAT1^75^, our findings for the first time provide evidence for a potential regulation of iPLA_2_β by NFκB and STAT1 at the transcriptional level.

Inhibition or genetic knockdown of iPLA_2_β decreases while overexpression of iPLA_2_β increases the susceptibility of islet β-cells to ER stress and apoptosis and reduces T1D incidence^4,11,12^. Here, we present evidence that both outcomes are impacted selectively by iPLA_2_β, but not iPLA_2_γ or cPLA_2_α. In view of this, we considered the impact of modulating iPLA_2_β on NFκB. We find that select chemical inhibition, siRNA-mediated downregulation, or CRISPR-Cas9-mediated reduction of iPLA_2_β reduces NFκB and STAT1 activation, suggesting their potential modulation by iPLA_2_β in ER stressed β-cells. When ChIP analyses were repeated with antibodies directed against iPLA_2_β, we find enrichment of iPLA_2_β at *Nf*κ*b* and *Stat1* promoter regions in ER-stressed β-cells, suggesting potential transcriptional regulation of *Nf*κ*b* and *Stat1* by iPLA_2_β. However, as iPLA_2_β does not contain DNA binding motifs, we considered the possibility that co-factors, previously not identified, facilitate such regulation.

Lipids are recognized to promote gene transcription^76–78^ and our lipidomics analyses reveal increased production of proinflammatory eicosanoids by β-cells with exposure to cytokines. Among the more prominent were PGE_2_, PGF_2_α, and 8-iso-PGF_2_α. These lipids induced β-cell apoptosis, that was associated with induction of NFκb, STAT1, and ER stress, suggesting that their signaling participates in ER stress-induced β-cell apoptosis. Interestingly, they also induced iPLA_2_β. This is not unprecedented as PGE_2_ has been reported to feedback regulate COX2^77^, which metabolizes arachidonic acid to generate PGs. In view of our findings, we assessed whether the PGs confer activity at the transcriptional level in ER stressed β-cells. Intriguingly, we find that the PGs promotes iPLA_2_β association with *Pla2g6* and *Nfkb* promoter regions, that were not significantly impacted in the presence of the iPLA_2_β inhibitor *S*-BEL, providing a likely mechanism by which the PGs serve as co-factors to promote association of iPLA_2_β with *Pla2g6* and *Nfkb* promoters.

This likely mechanism, however, was not congruent with the findings that chemical inhibition of iPLA_2_β, which would be expected to reduce eicosanoid production^79^, reduced NFκB and STAT1 activation, ER stress, and β-cell apoptosis, though it did not impair iPLA_2_β enrichment at *Pla2g6* or *Nf*κ*b* promoter regions. Further assessments with chemical inhibitors revealed that only inhibition of iPLA_2_β, but not of iPLA_2_β or cPLA_2_α, reduced iPLA_2_β. Taken together, we posit that there are two separate but required processes in play: one, is the cytokine-induced emrichment of iPLA_2_β with promoter regions, which does not require co-factors. Second and subsequent to iPLA_2_β association with promoters, triggering of its transcriptional “activity” is facilitated by the tested PGs, serving as co-factors/co-activators. This is consistent with the mitigated ER stress outcomes with chemical inhibition of iPLA_2_β, though its association with promoters remains intact. Such a mechanism may also explain why under basal conditions, overexpression of iPLA_2_β does not induce deleterious outcomes, but under stress they are unmasked^4,80^. This is consistent with a prerequisite need for *de novo* generated lipid signaling to facilitate iPLA_2_β activity at the promoter regions and warrants further investigation.

Our findings provide a rational novel sequence of feedback events (**Fig.13**), where ER stress in β-cells (a) induces NFκB and STAT1, which in turn induce *Pla2g6* to increase expression iPLA_2_β and (b) induces iPLA_2_β localization to nucleus and its association with *Pla2g6*, *Nf*κ*b,* and *Stat1* promoter regions, and (c) leads to increased generation of iPLA_2_β- derived PGs which facilitate iPLA_2_β transcriptional activity. This sequence of events links select lipid signaling with triggering of transcription of NFκB and STAT1 genes, leading to generation of apoptosis-related factors that contribute to β-cell death. Importantly, both cytokines and the select iDLs promoted association of iPLA_2_β with *Pla2g6* and *Nf*κ*b* promoter regions suggesting that ER stress elicits the same sequence of events, in the context of human islet β-cell apoptosis leading to T1D development.

## Author’s Contributions

**XL** was involved with the design, experimentation, data analyses and manuscript preparation; **AKC** generated the CRISPR-Cas9 guides, **TA** contributed to NFκB immunoblotting; **SEN** contributed to the design of the ChIP experiments; **DS**, **AF**, and **CEC** performed the lipidomics mass spectrometry and data analyses; **CSH**, **ARW**, and **ESN** contributed to the transcriptional approaches; **YGT** assisted with the cell culturing and human islets experiments; and **SR** oversaw all experimental design, data analyses and preparation of the manuscript. All co-authors contributed to editing of the manuscript.

## Acknowledgements

This work was supported in part by funding (to SR) from NIH/NIDDK (R01DK069455, R01DK110292), NIH/HIRN (U01 DK127786), Breakthrough T1D (3-SRA-2023-1362-S-B), Beatson Foundation (#2023-20), UAB-Department of Cell Developmental, and Integrative Biology, UAB-Comprehensive Diabetes Center, and UAB Diabetes Research Center (P30 DK079626). The contents of this manuscript do not represent the views of the Department of Veterans Affairs or the United States Government. This work was mainly supported by the National Institutes of Health via R01DK126444 (to CEC, SR), the Veteran’s Administration (VA Merit Review, BX001792 (to CEC), VA Merit Review award, BX006063 (to CEC), VA Merit Review award, RD001334 (to CEC), and a Senior Research Career Scientist Award, IK6RD002739 (to CEC)). This work was also peripherally supported by way of research cores, methodology and technology development, and software/bioinformatic development by the National Institutes of Health via P01 CA171983 (to CEC), P01 CA302570 (to CEC), NIH/NCI Cancer Center Support Grant P30 CA044579, R01 AI139072 (to CEC), and R01 GM155691 (to CEC). We thank Dr. George Kokotos, University of Athens for the generous supply of FKGK18. Activity of the AIDP is approved by the Health Research Ethics Board at the University of Alberta (Approval # Pro-00001620).

## Supplemental Figures

**Supplemental Figure 1.**
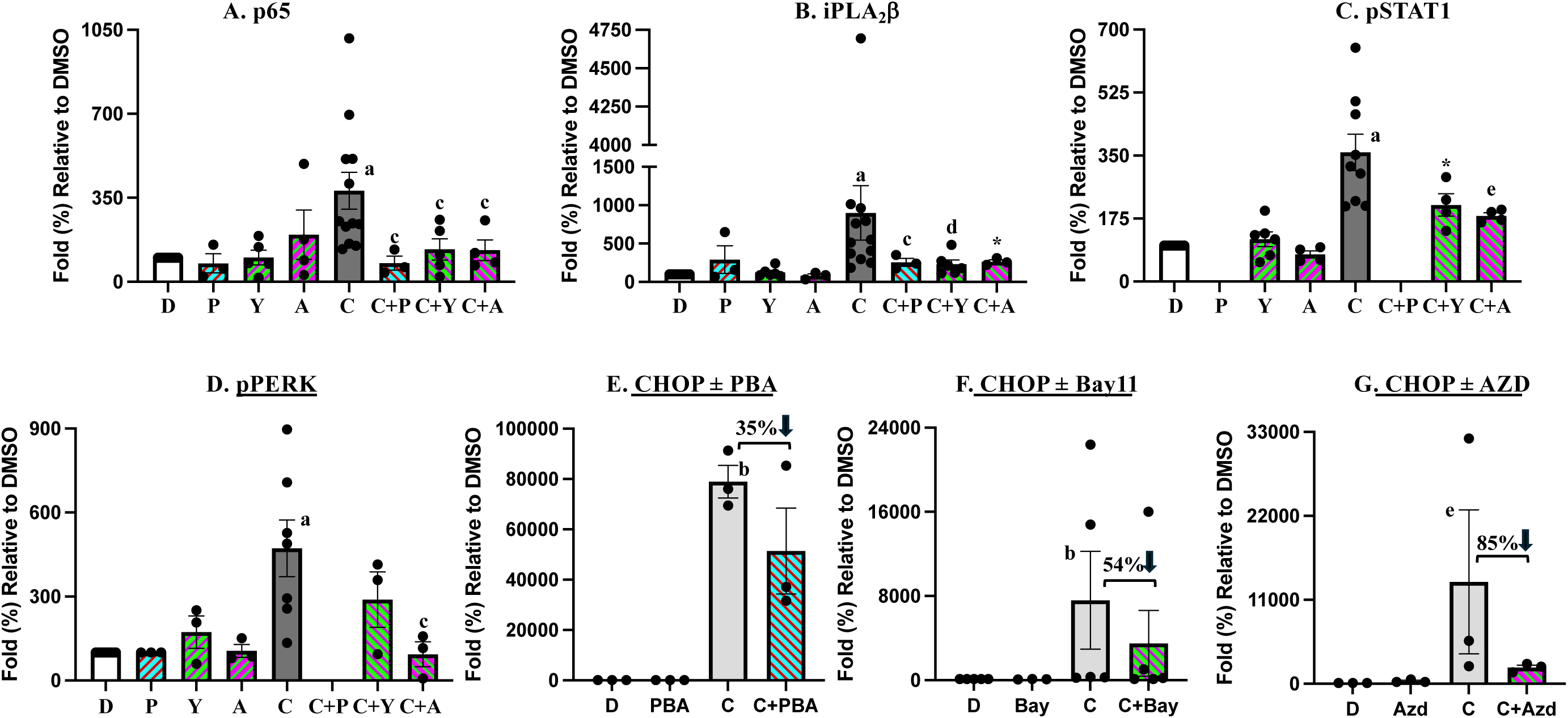
Impact of ER Stress, NFκB, and STAT1 on iPLA_2_β and Stress Factors. MIN-6 β-cells were treated with cytokines, as in **Figs. 1 and 2** in the absence or presence of <u>P</u>BA, BA<u>Y</u> 11, or <u>A</u>ZD for 16h. Cytosols were then prepared and processed for immunoblotting analyses for p65-NFκb, iPLA_2_β, pSTAT1, pPERK, and CHOP. (Cytokine groups significantly different from corresponding DMSO groups, ^a^p < 0.0001 and _b_p = 0.0003; ^c,d,*,e^Cytokine + intervention groups significantly different from corresponding cytokine groups, p < 0.05, p < 0.01, p < 0.05 (1-tailed), and p < 0.005, respectively.

**Supplemental Figure 2.**
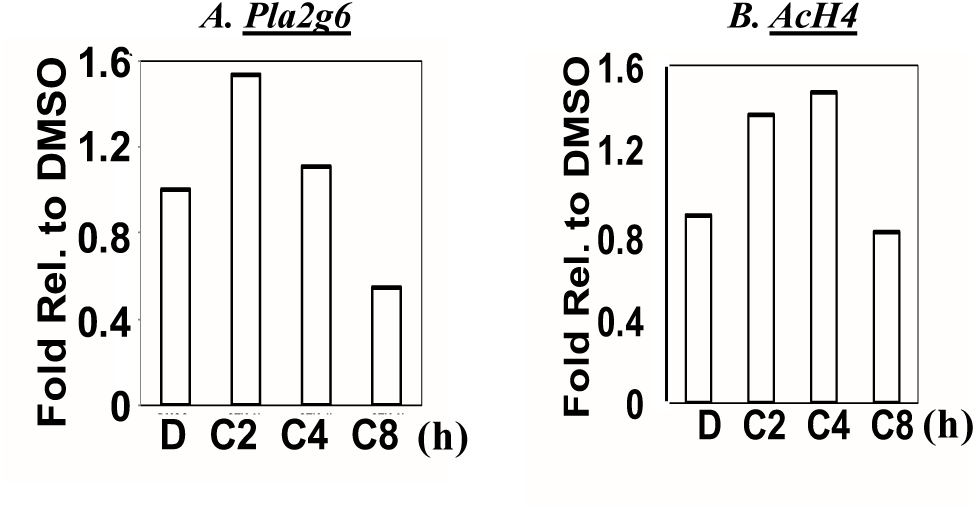
Time-Course of iPLA_2_β Association with *Pla2g6* Promoter Region. INS-1 β-cells were treated with <u>D</u>MSO or <u>C</u>ytokines, as in **Suppl. Fig. 1**, for 2-8h and then processed for ChIP analyses using select antibodies for iPLA_2_β and AcH4 and subsequent qPCR analyses for *Pla2g6* promoter region.

**Supplemental Figure 3.**
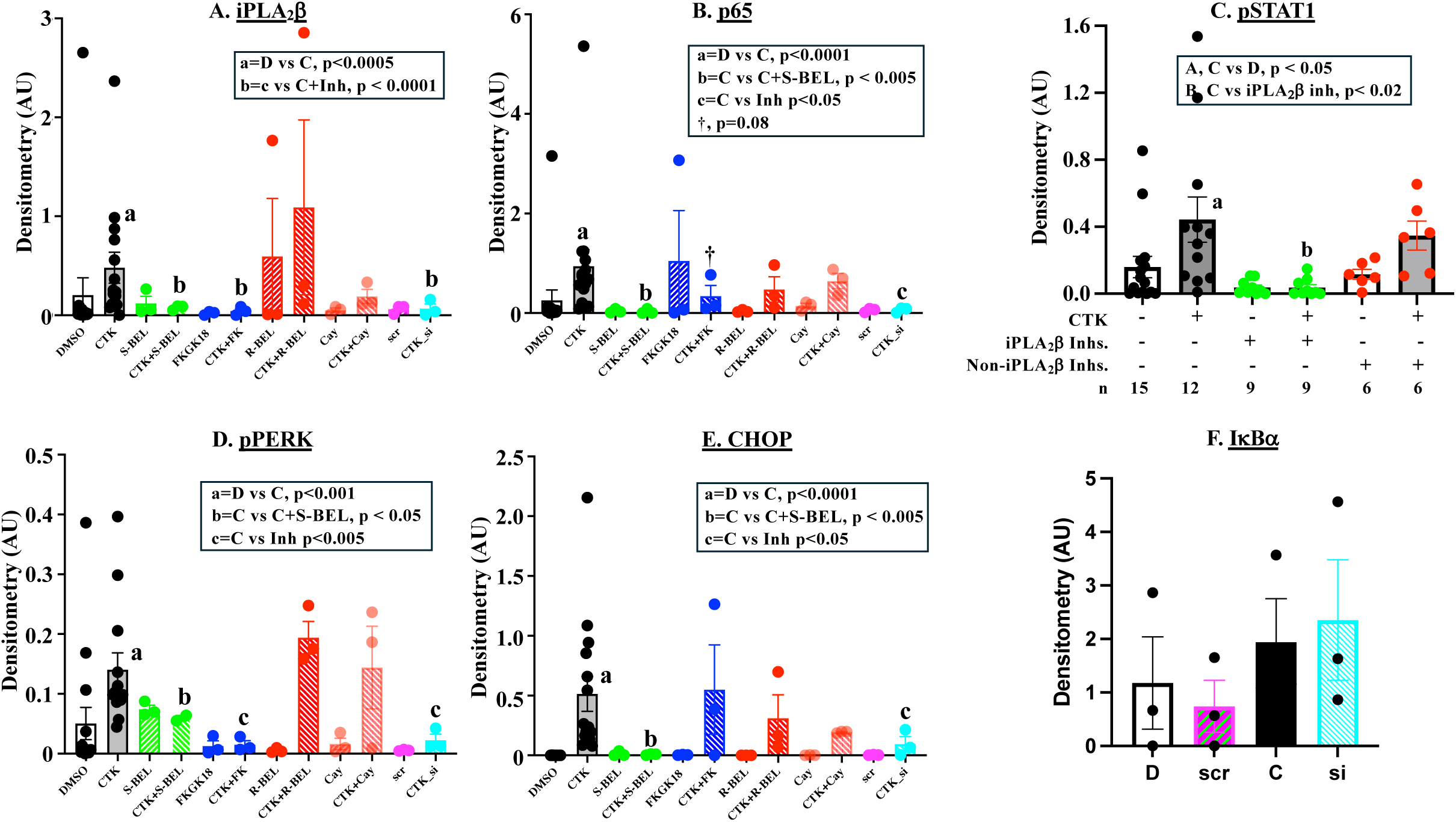
Select Targeting of iPLA_2_β Reduces Stress Responses. MIN-6 β-cells were treated with <u>D</u>MSO or cytokines for 24h, as in **Suppl. Fig. 1**, in the absence or presence of <u>D</u>MSO, <u>P</u>BA, *S*-BEL, <u>F</u>KGK18, *<u>R</u>*-BEL, <u>C</u>ay10502, <u>scr</u>ambled RNA, or <u>si</u>RNA targeting *Pla2g6*. Cytosols were then prepared and processed for immunoblotting analyses for p65-NFκB, iPLA_2_β, pSTAT1, pPERK, and CHOP, and IκBα immunoblotting analyses. (Significance differences are presented as inserts.)

**Supplemental Figure 4.**
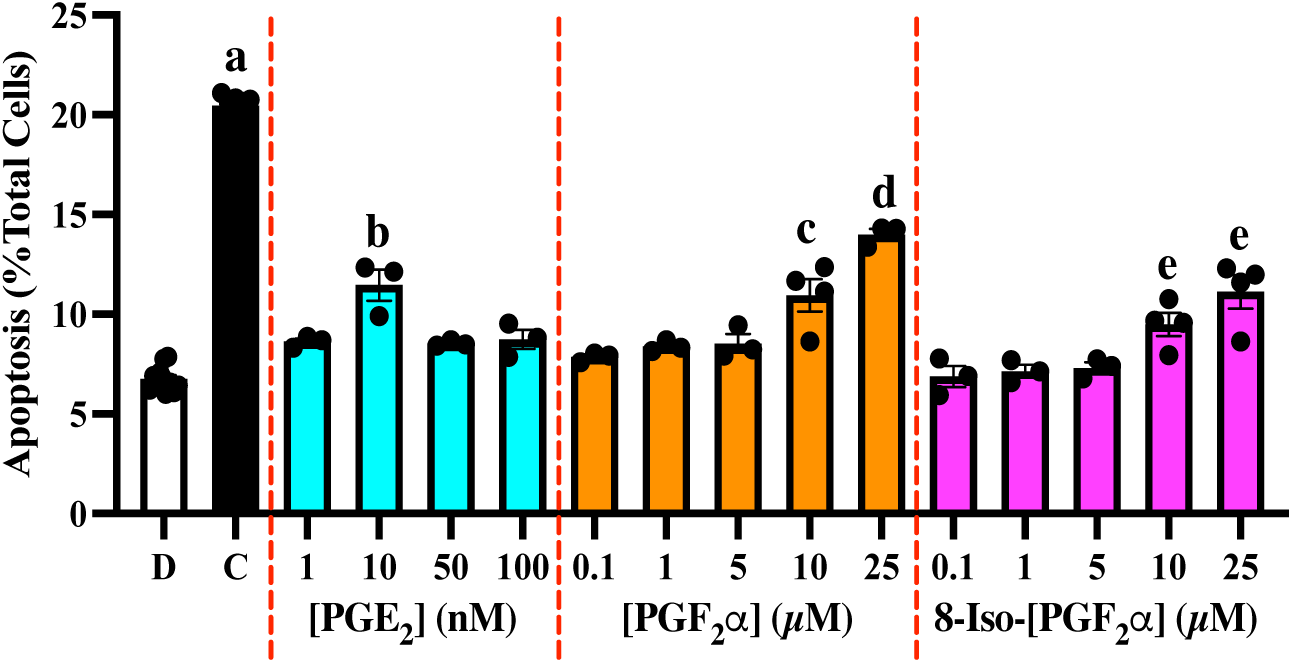
Concentration-Dependent Induction of β-cells by Select Prostaglandins. MIN-6 β-cells were treated with <u>D</u>MSO, <u>C</u>ytokines, or PGs (10 nM PGE_2_, 10 µM PGF_2_α, 10 µM 8-Iso-PGF_2_α) for 24h and processed for TUNEL. The mean ± SEMs of %apoptotic cells, relative to total cells, are presented. (^a-^ _e_Significantly different from DMSO group, p < 0.05 - p <0.001.)

**Supplemental Figure 5.**
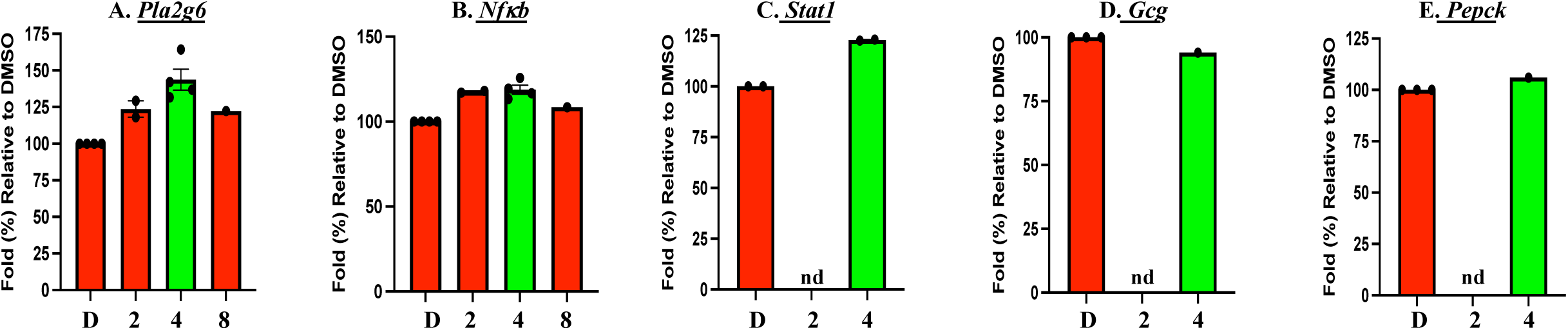
Time-Course of Select PGs-Induced Enrichment of iPLA_2_β at Promoter Regions. MIN6 β-cells were treated with <u>D</u>MSO or <u>PGs</u> cocktail (10 nM PGE_2_ + 10 µM PGF_2_α + 10 µM 8-Iso-PGF_2_α) for 2-8h. The cells were then processed for ChIP analyses using select antibodies for iPLA_2_β, and subsequently for qPCR analyses of *Pla2g6, Nfkb, Stat1, Gcg,* and *Pepck* promoter regions.

**Supplemental Figure 6.**
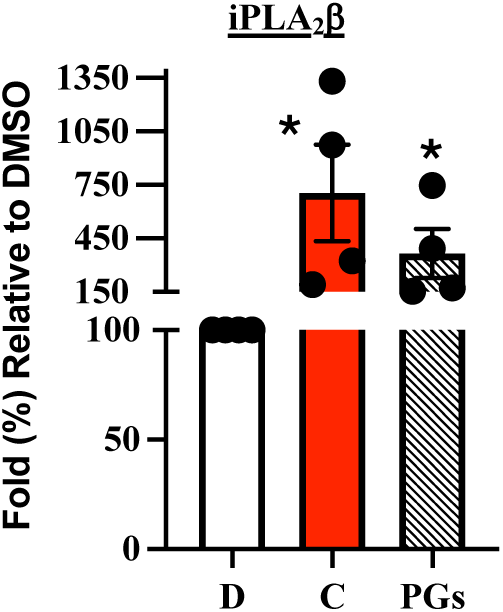
Impact of Select PGs on iPLA_2_β Expression. MIN6 β-cells were treated with <u>D</u>MSO or <u>PGs</u> cocktail for 16h and then processed for iPLA_2_β immunoblotting analyses. (^†^Significantly different from D, p < 0.05, 1-tailed test, n=4/group.)

**Supplemental Figure 7.**
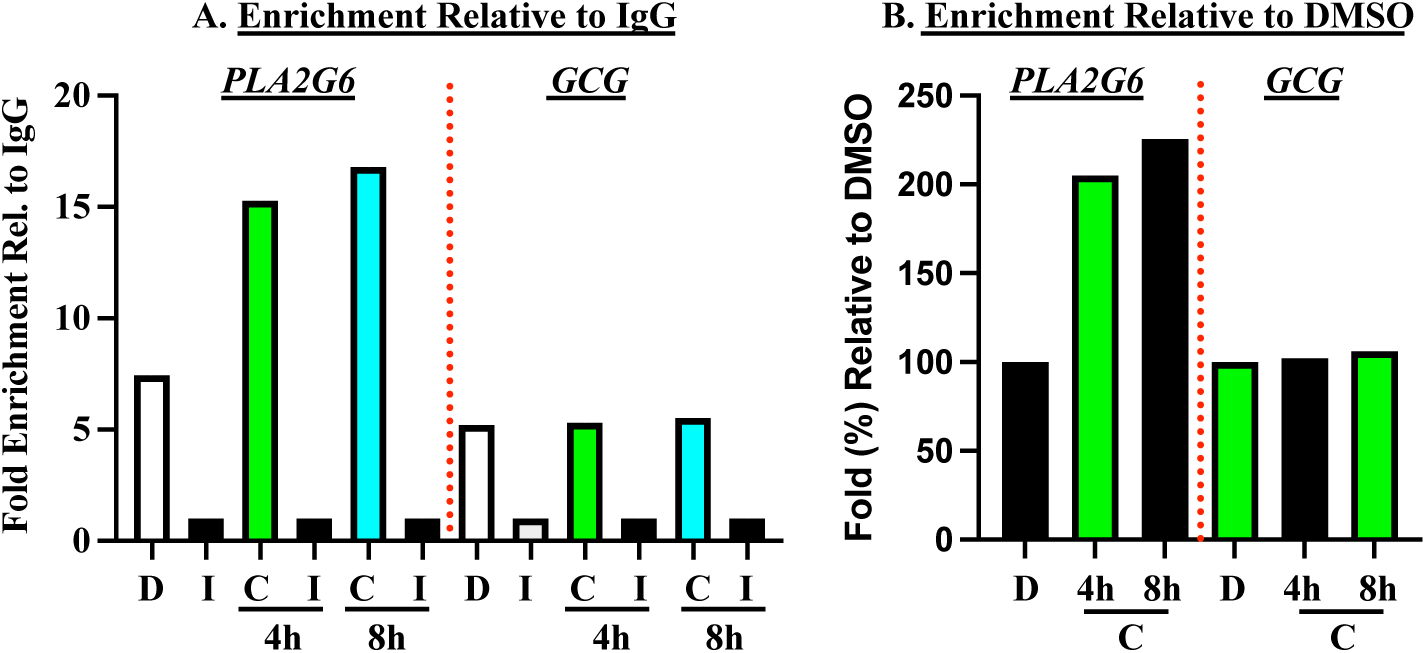
Time-Course of Cytokines-Induced Enrichment of p65-NFKB at Promoter Regions in Human Islets. Human islets from health donors (3500-5000) were treated with <u>D</u>MSO or <u>C</u>ytokines (100 U/ml IL-1β + 300 U/ml IFNγ) for 4 and 8h. The islets were then processed for ChIP analyses using select antibodies for p65-NFkB and subsequent qPCR analyses for *PLA2G6* and *GcG* promoter regions. Enrichment relative to IgG (**A**) and DMSO (**B**) are presented.

